# Anterior cingulate cortex in complex associative learning: monitoring action state and action content

**DOI:** 10.1101/2025.01.29.635442

**Authors:** Wenqiang Huang, Arron F Hall, Natalia Kawalec, Ashley N Opalka, Jun Liu, Dong V Wang

## Abstract

Environmental changes necessitate adaptive responses, and thus the ability to monitor one’s actions and their connection to specific cues and outcomes is crucial for survival. The anterior cingulate cortex (ACC) is implicated in these processes, yet its precise role in action monitoring versus outcome tracking remains unclear. To investigate this, we developed a novel discrimination–avoidance task for mice, designed with clear temporal separation between actions and outcomes. Our findings show that ACC neurons primarily encode post-action variables over extended periods, reflecting the animal’s preceding actions rather than the outcomes or values of those actions. Specifically, we identified two distinct subpopulations of ACC neurons: one encoding the action state (whether an action was taken) and the other encoding the action content (which action was taken). Importantly, increased post-action ACC activity was associated with better performance in subsequent trials. These findings suggest that the ACC supports complex associative learning through extended signaling of rich action-relevant information, thereby bridging cue, action, and outcome associations.

## Introduction

The ability to adapt behavior based on environmental cues and outcome-related feedback is essential for survival. Central to this process is the brain’s capacity to integrate diverse information to form cue–action–outcome associations that guide future behaviors across varying conditions. However, the neural mechanisms underlying this cognitive flexibility remain poorly understood. The anterior cingulate cortex (ACC) has emerged as a critical node in mediating this process, with growing evidence underscoring its pivotal role in higher-order cognition and adaptive behavior [1–7]. Despite this, the precise role of the ACC in updating and modifying behavior is still under debate [1–7].

Historically, the ACC was thought to be essential for error detection or conflict monitoring [8–10]. However, accumulating evidence has challenged these views [11–17], leading to proposals that the ACC may support a diverse range of functions. These include value encoding, strategy updating, decision-making, action monitoring, and outcome tracking [18–29]. Some of these proposed functions remain controversial, such as the ACC’s role in decision-making [18–20], while others are defined more broadly, such as the ACC’s role in strategy updating [26, 28]. Nevertheless, there is broad agreement that ACC activity is closely linked to actions and/or action-related outcomes, particularly in tasks involving competing actions [18–29]. Supporting this view, lesions of the ACC impair the maintenance of newly acquired task performance and disrupt behavioral flexibility, including the ability to associate actions with their outcomes [27–30].

Despite evidence linking the ACC to action-related cognitive processes, past studies have primarily relied on simple motor tasks, such as saccades, licking, or joystick movements, leaving its role in more naturalistic behaviors largely unexplored. Moreover, the brief temporal separations between cues, actions, and outcomes in prior studies have made it challenging to determine whether ACC neuronal activity contributes to decision-making, action monitoring, or merely tracking the outcomes of those actions [17–31]. In this study, we aim to disentangle the ACC’s role by employing a novel discrimination–avoidance task, designed to evoke robust, naturalistic action responses and establish a clear temporal separation between cues, actions, and outcomes. Our findings highlight a distinct role of the ACC in encoding post-action variables that capture detailed information about preceding actions, rather than tracking the outcomes of those actions or their associated values.

## Results

### A novel discrimination–avoidance task

We first developed a novel discrimination–avoidance task, tailored to investigate action-based complex associative learning in mice. In this task, animals learn to discriminate between two auditory cues that predict context-dependent footshocks. Specifically, sounds A and B signal electric shocks in rooms A and B of a shuttle box at sound terminations, respectively (Fig. 1A; Fig. S1). This design requires animals to either “stay” in the current room or “shuttle” to the adjacent room during sound presentations to avoid shocks (Supplementary Videos 1–2). During inter-trial intervals, animals are free to explore either room without consequence.

**Figure 1.**
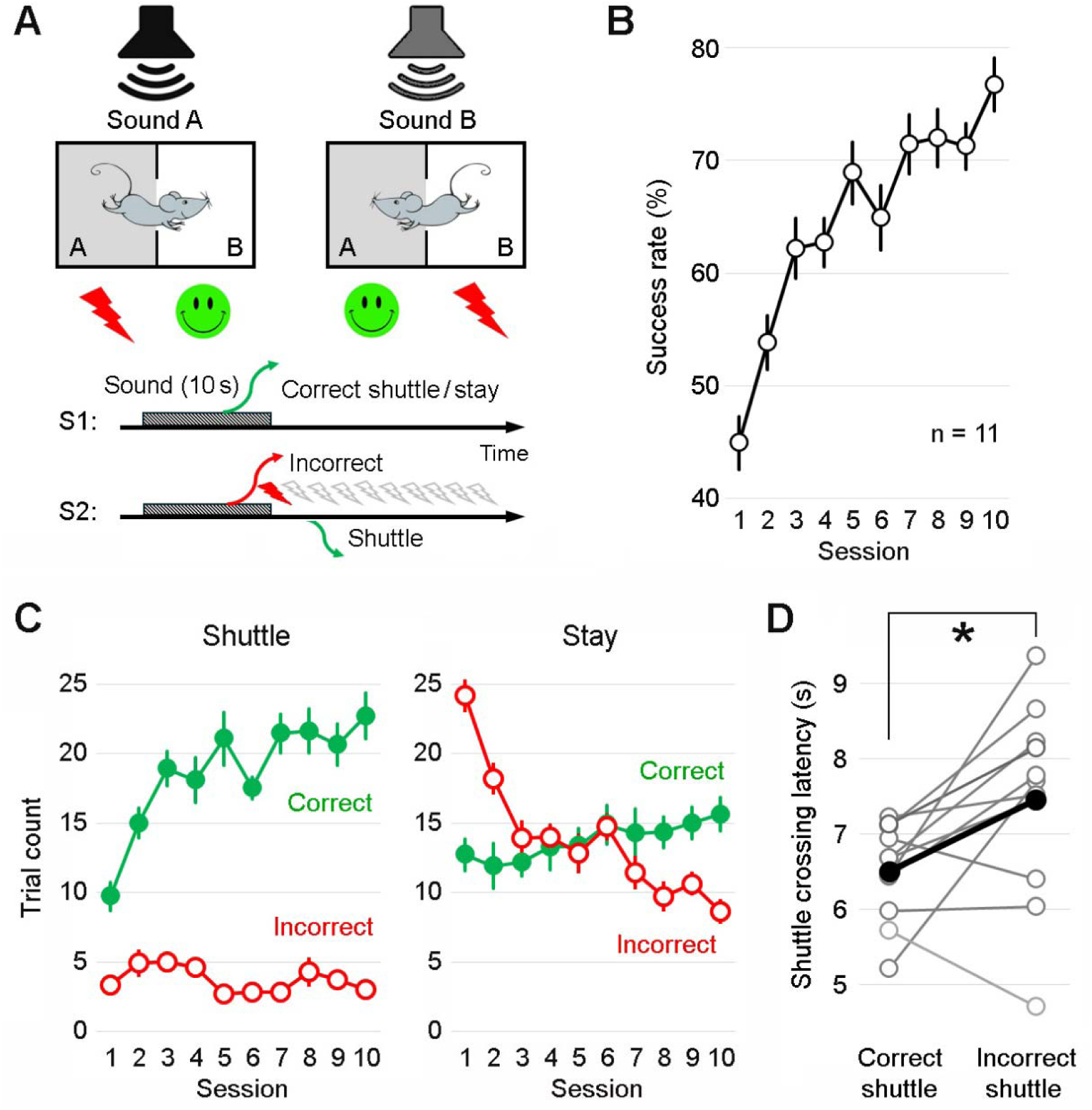
A novel discrimination–avoidance task. **A,** Top: Schematic of the task. Mice are trained to discriminate between two auditory cues (lasting 10 s) and shuttle between two distinct rooms of a shuttle box to avoid footshocks. Specifically, sounds A and B signal shocks in rooms A and B, respectively. Bottom: Two behavioral response scenarios. S1: If the mouse makes the correct response, either by staying in or shuttling to the correct room before the sound ends, no shock is administered. S2: If the mouse makes an incorrect response, either by staying in or shuttling to the incorrect room before the sound ends, up to 10 mild shocks (0.5 mA, 0.1 s; 2 s apart) are administered until the mouse shuttles to the correct room. Each training session comprises 50 trials, 60 s apart; sounds A and B are presented in a pseudorandom order. **B,** Learning curve showing the success rate in avoiding shocks across training sessions (mean ± s.e.m.; n = 11 mice). F_9, 90_ = 14.78; P < 0.001; One-way ANOVA. **C,** Correct and incorrect trial counts for shuttle *vs.* stay trials across training sessions for the same mice shown in B. Correct shuttle: F_9, 90_ = 14.62, P < 0.001; Incorrect shuttle: F_9, 90_ = 1.94, P = 0.057; Correct stay: F_9, 90_ = 1.55, P = 0.142; Incorrect stay: F_9, 90_ = 18.73, P < 0.001; One-way ANOVA. **D,** The mean shuttle crossing latency, averaged over the last two sessions for individual mice, is significantly shorter in correct trials than that in incorrect trials (t_10_ = 2.62, P = 0.026; Effect Size: Cohen’s d = 0.789; Power = 0.656; paired *t* test). The black line indicates the mean; grey lines indicate individual mice. Shuttle crossing is defined as the body center crossing the midline opening of the shuttle box.

Notably, the auditory cues are not inherently associated with positive or negative valence; instead, their meaning is dynamically determined by the animal’s current location (room A or B) at the time of cue presentation, signaling either safety or shock. Thus, this task, which requires discrimination of sensory cues and environmental contexts, as well as the integration of cues, actions, and outcomes, serves as an ideal tool to study complex associative learning. Our results showed that mice gradually learned the task, achieving an average success rate of 76.7% in avoiding shocks by the 10th training session (Fig. 1B). An additional five training sessions yielded a modest improvement, increasing the average success rate to 82.5% on the 15th session (Fig. S2).

Given that the task requires two distinct behavioral responses, we separated the trials into shuttle and stay trials for further analysis. In shuttle trials, mice must move to the adjacent room before the sound ends to avoid shocks. Across training sessions, correct shuttles increased markedly, whereas incorrect shuttles decreased only modestly (Fig. 1C; Fig. S2). This pattern suggests that animals choose to shuttle primarily when they are confident in the outcome (i.e., safety); otherwise, they remain in place. Consistent with this notion, in stay trials, where mice must remain in their current room to avoid shocks, incorrect stays decreased markedly across training sessions, mirroring the improvement in correct shuttle performance. By contrast, correct stays increased only modestly (Fig. 1C; Fig. S2).

We also compared shuttle response latencies between correct and incorrect trials during the late stages of training. On average, the shuttle response latency was 6.5 s during correct shuttles, providing a 3.5-s temporal separation between actions (shuttles) and outcomes (safety or shocks; Fig. 1D). This clear temporal distinction allows us to compare how information is coded during the action *vs.* outcome periods. The longer shuttle latency during incorrect shuttles suggests that last-second responses are more likely to be incorrect (Fig. 1D).

### ACC neurons exhibit robust post-action firing changes

Next, we conducted multi-channel *in vivo* electrophysiology recordings from the ACC (Fig. 2A; Fig. S2) in mice performing the discrimination–avoidance task after they had completed 10 training sessions and achieved a success rate of 70% or above. We initially observed robust changes in ACC activity following shuttle responses (Fig. 2 B&C). While many ACC neurons showed changes in activity during the shuttle period, the majority of these changes persisted well after shuttle termination, with some lasting up to 30 s after the initial response (Fig. 2 C&E; Fig. S2). In contrast, the same ACC neurons showed no or limited activity changes during stay trials in response to the auditory cues (Fig. 2 D&F).

**Figure 2.**
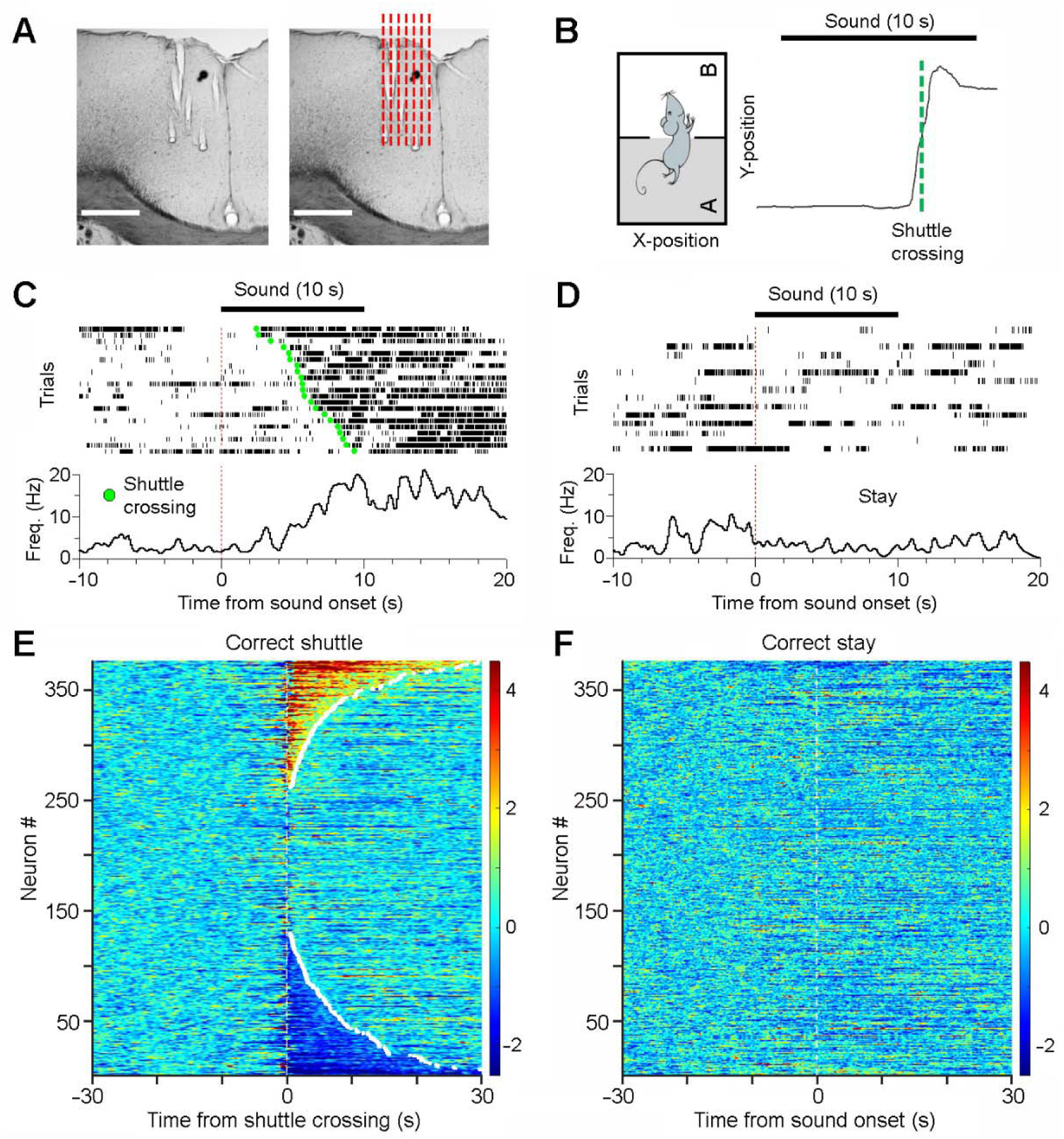
ACC neurons primarily encode post-action variables. **A**, A representative brain section showing electrode tracks (left) and the presumed implantation sites (red dashed lines; right), located largely within the ACC. Scale bars, 0.5 mm. **B,** A representative shuttle response and corresponding Y-position of the animal’s body center. **C&D,** Peri-event rasters and histograms of a representative ACC neuron during correct-shuttle (C; trials sorted by shuttle latency) and correct-stay trials (D). In this session, there were 21 correct shuttles, 15 correct stays, 5 incorrect shuttles, and 9 incorrect stays. **E&F,** Heatmaps showing the activity of all recorded ACC neurons (n = 376) during correct-shuttle (E) and correct-stay trials (F). Neurons in E and F are arranged in the same order.

To assess ACC neuronal population activity dynamics, we performed Principal Component Analysis (PCA) on simultaneously recorded ACC neurons across correct-shuttle and correct-stay trials (Fig. 3A). The first three principal components (PCs) revealed pronounced changes in population activity during shuttle trials, whereas activity remained relatively stable during stay trials (Fig. 3B; Fig. S3). Quantifying the maximum trajectory distance within the 3D PC space revealed consistent, prominent changes in population activity during shuttle responses across animals in five representative sessions (Fig. 3C). The pronounced ACC activity during and following shuttles, coupled with minimal activity in response to cues, suggests that ACC neurons primarily encode action and post-action variables. Such sustained post-action activity may support complex associative learning by linking temporally separated action- and outcome-relevant information.

**Figure 3.**
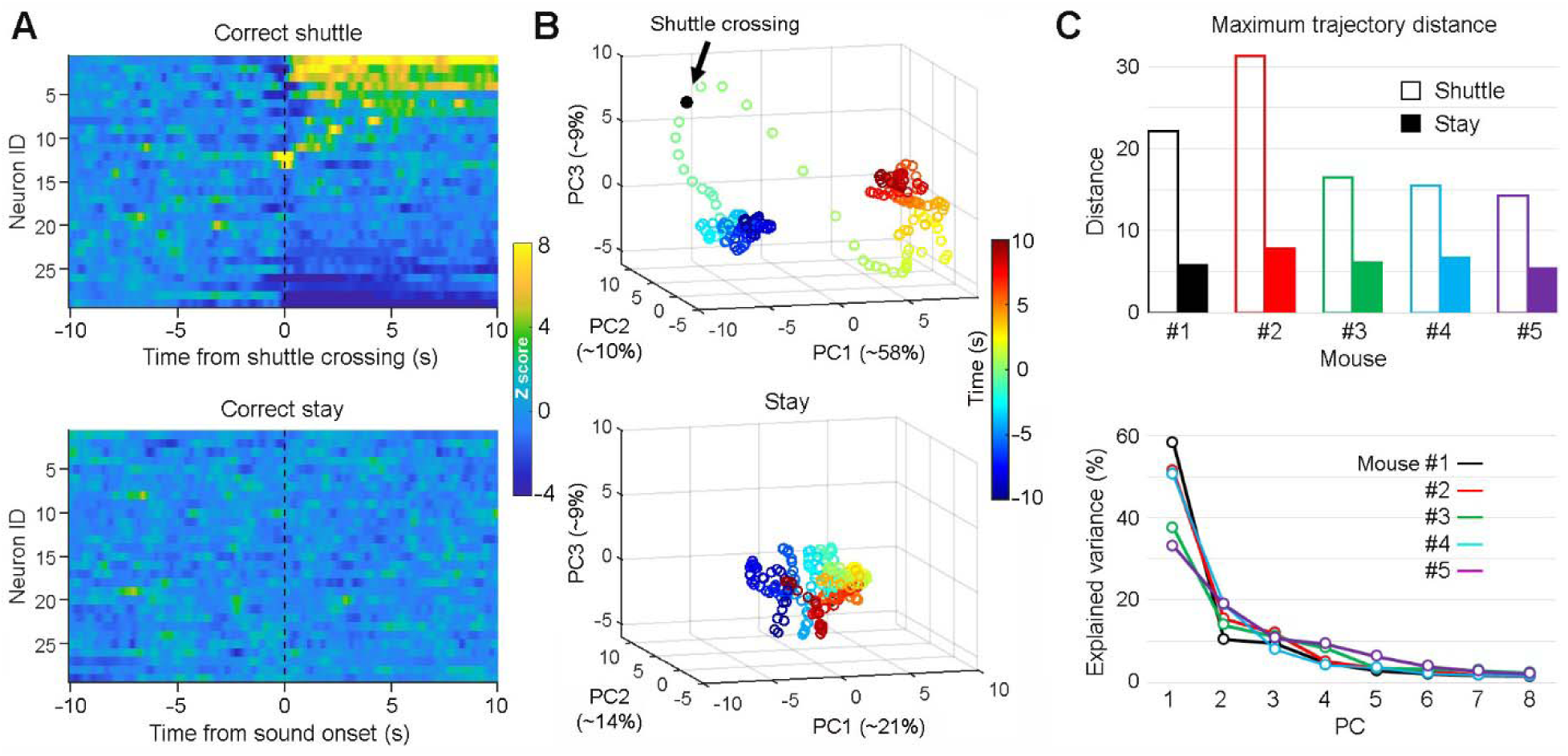
ACC neurons primarily respond during “shuttle” but not “stay” trials. **A,** Heatmaps showing the activity of simultaneously recorded ACC neurons (n = 29) during correct-shuttle (top) and correct-stay trials (bottom) from the same recording session as shown in Fig. 2C. **B,** Principal component analysis (PCA) of ACC neuronal population activity as shown in A. PC1, PC2, and PC3 are the first three principal components; the numbers are the percentages of total variance explained by the corresponding PCs. Each circle indicates a time lapse of 0.1 s. Note that there is a robust neural state change in the 3-D PC space surrounding the shuttle response (top), but not the stay response (bottom). **C,** Maximum trajectory distance between the center of the baseline period and any post–Time 0 data point, as shown in B, across five animals (top), and the corresponding scree plots showing variance explained by the first eight PCs during shuttle trials (bottom).

### Temporal organization of ACC activity during shuttle behavior

To assess whether ACC activity has a role in encoding pre-action variables, we examined ACC activity aligned to shuttle initiations, defined as locomotion velocity exceeding one standard deviation (s.d.) above baseline (Fig. 4A). Our results revealed diverse response properties of the ACC neuronal population (Fig. 4B), which frequently persisted beyond the shuttle response window (∼1–3 s between shuttle initiations and terminations; Fig. 4A). PCA classified these responses into three major categories (Fig. 4 C&D). Pre-shuttle ramping activity was mainly observed in a small subset of neurons, (Types 3 b&c, ∼16% of the population), indicating a limited role of the ACC in pre-action information coding, including decision-making and planning processes. The remaining ACC neurons primarily displayed sustained activity after initiation, either activation (Types 1a, 1b, 3a) or inhibition (Types 2a, 2b), highlighting the predominant role of the ACC in post-action information processing. Overall, despite aligning to shuttle crossings or initiations, the extended analysis windows largely captured post-action ACC activity.

**Fig. 4.**
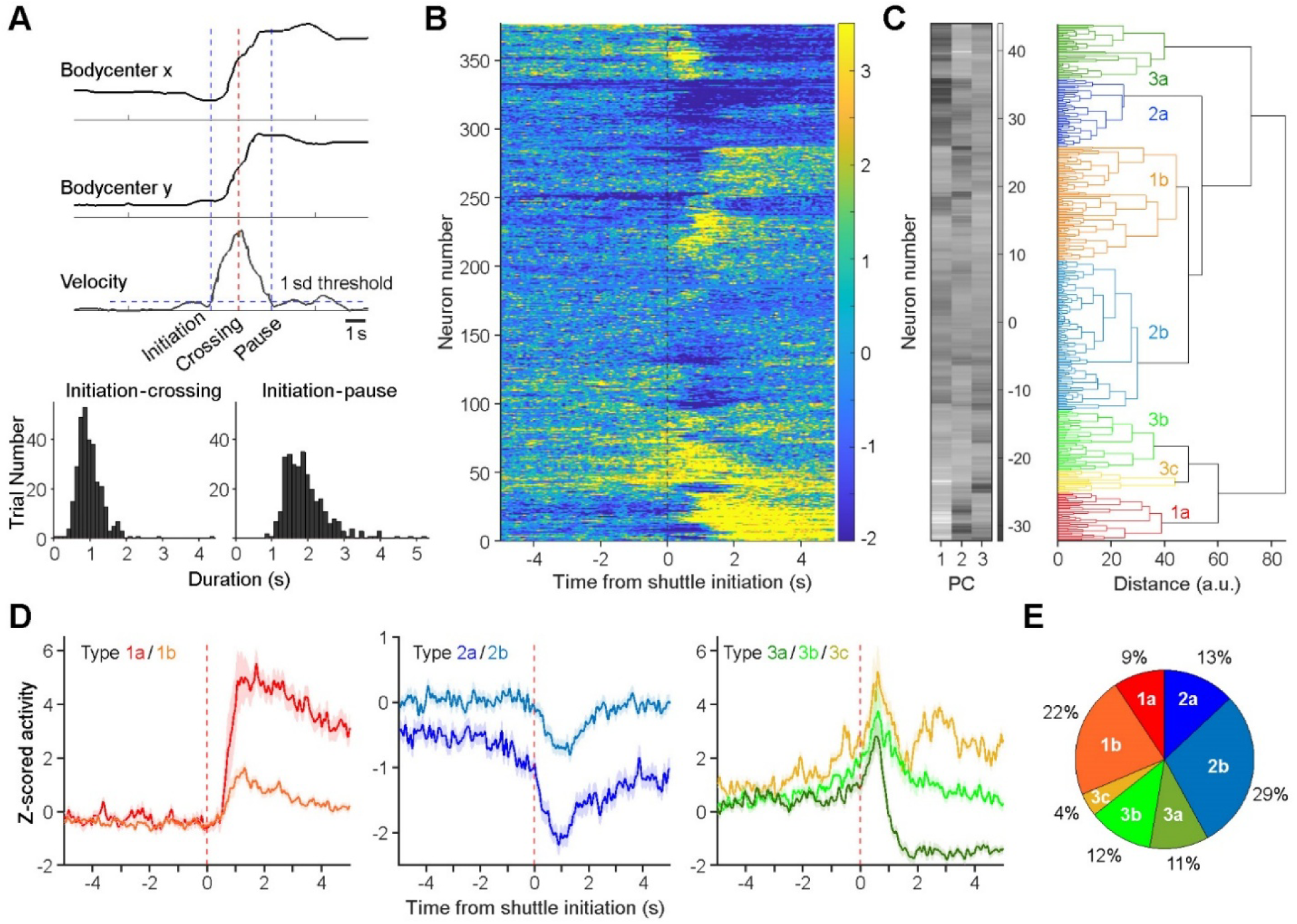
Characterizing ACC neuronal activity in relation to action initiations. **A**, Top, schematic illustrating the definition of shuttle initiation, crossing, and termination (pause). Bottom, distributions of half- (left) and full-shuttle durations (right). **B**, Z-scored activity of all ACC neurons (n = 376) during correct shuttle trials. **C,** Principal-component analysis (PCA) classifies ACC neuronal activity (as shown in B) into seven categories. PC1, PC2, and PC3 represent the first three principal components color coded from low (dark) to high scores (white). **D**, Mean activity (± s.e.m.) of the seven categories of ACC neurons. **E**, Fractions of individual categories of ACC neurons.

To determine which shuttle event (initiation, crossing, or termination) captured the most acute changes in ACC neuronal firing, we conducted an event-locked modulation analysis (Fig. S4). Our results showed that shuttle crossing was associated with the largest fraction of significantly modulated ACC neurons, particularly when comparing short windows (250–1000 ms) between pre- and post-event ACC activity (Fig. S4). These findings suggest that shuttle crossing represents the most prominent event for ACC engagement during shuttle behaviors.

### ACC neurons exhibit limited modulation by speed

Given that the post-action ACC activity is often prolonged and outlasts shuttle termination, we hypothesized that this activity is distinct from locomotion encoding. To test this, we analyzed ACC activity in relation to movement speed. Each task session included a 5-min free exploration period before trials began, allowing us to assess if ACC activity is associated with basic motor functions. We found that only a small portion of neurons showed speed-correlated activity, with either positive (7.7%) or negative correlation (6.9%), indicating that the majority of the neurons are not tuned to movement speed (Fig. 5 A–C). This is consistent with our observation of sustained post-shuttle ACC activity in the absence of movement, which is distinct from locomotion encoding. Nevertheless, it remains unclear whether this small fraction of speed-related neurons represents a distinct subpopulation within the ACC or reflects recordings from nearby motor cortex. Postmortem examination of the recording sites suggests that most neurons were recorded from the ACC, while a small subset was located at the border between the ACC and motor cortex (Fig. S2). Therefore, it is possible that the speed-related neurons originated from the motor cortex.

**Figure 5.**
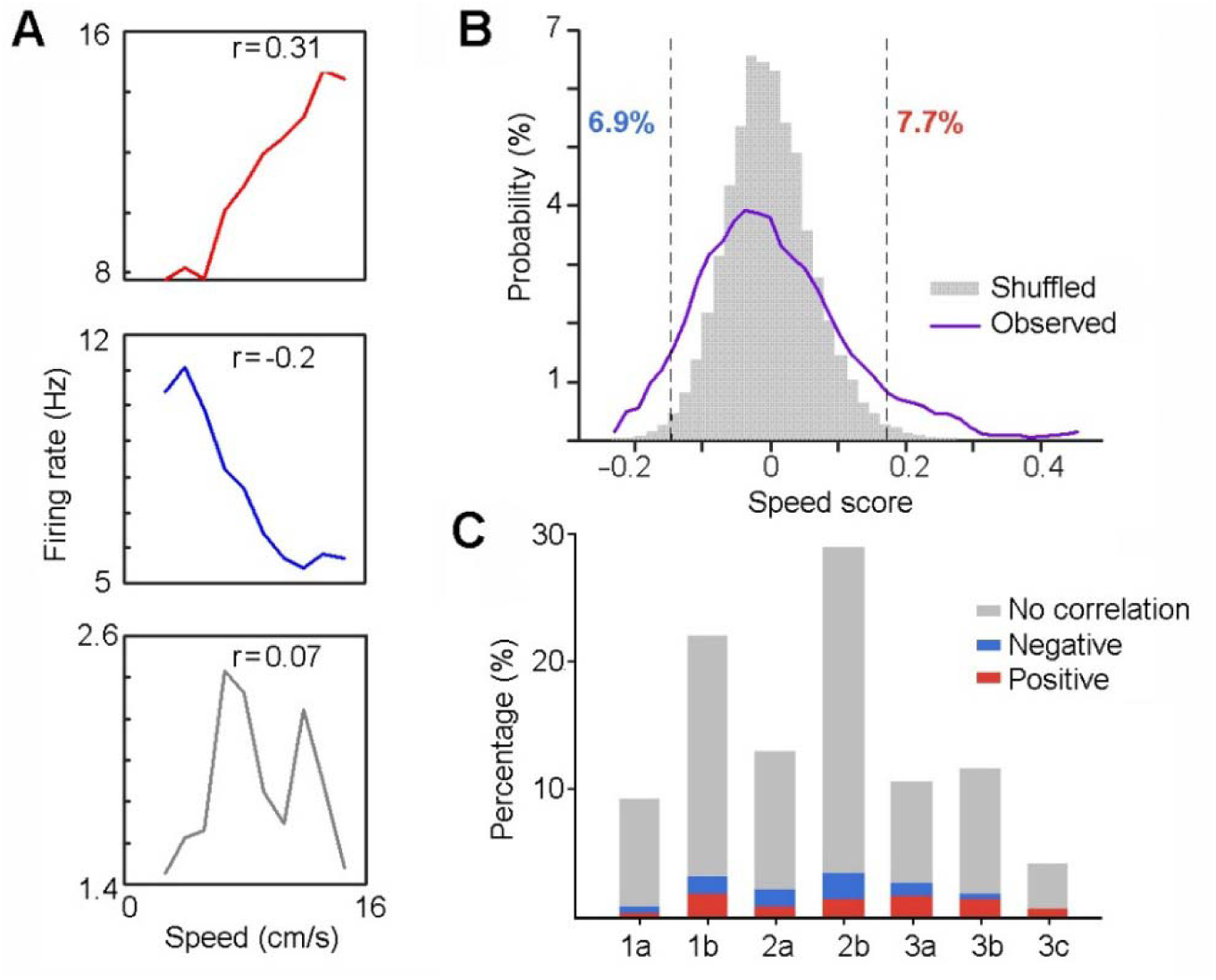
ACC neurons exhibit limited modulation by speed. **A,** Speed-tuning curves of three simultaneously recorded neurons during free exploration in shuttle boxes, showing positive (red), negative (blue), or no correlation (gray) between firing rate and locomotion speed. “r” indicates Pearson’s correlation coefficient. **B,** Distributions of observed *vs.* shuffled correlation values across all recorded ACC neurons (n = 376). Dashed lines mark the 99th percentile of shuffled distribution. Overall, 7.7% of recorded neurons were positively modulated by speed, and 6.9% were negatively modulated. **C,** Proportions of speed-modulation for each category of ACC neurons (see Fig. 4D).

### ACC neurons monitor actions independent of outcomes

To determine if the post-action ACC activity encodes information related to outcomes, we analyzed ACC neuronal activity across three distinct conditions: correct shuttles, incorrect shuttles, and post-shock shuttles (Fig. 6A; post-shock shuttles are defined as shuttles following footshocks received during incorrect-shuttle and incorrect-stay trials). Our results revealed that ACC neurons exhibited similar activity patterns across all three conditions, regardless of outcomes (presumed safety *vs.* uncertainty *vs.* safety; Fig. 6B). Specifically, both activity strength (Fig. 6C) and activity pattern (Fig. 6D) were significantly correlated across conditions, indicating outcome-independent responses in ACC neurons. Consistently, we found that ACC neurons showed limited responses to footshocks during incorrect trials (Fig. S5). Together, these findings suggest that ACC neurons monitor actions independent of outcomes or values associated with these actions.

**Figure 6.**
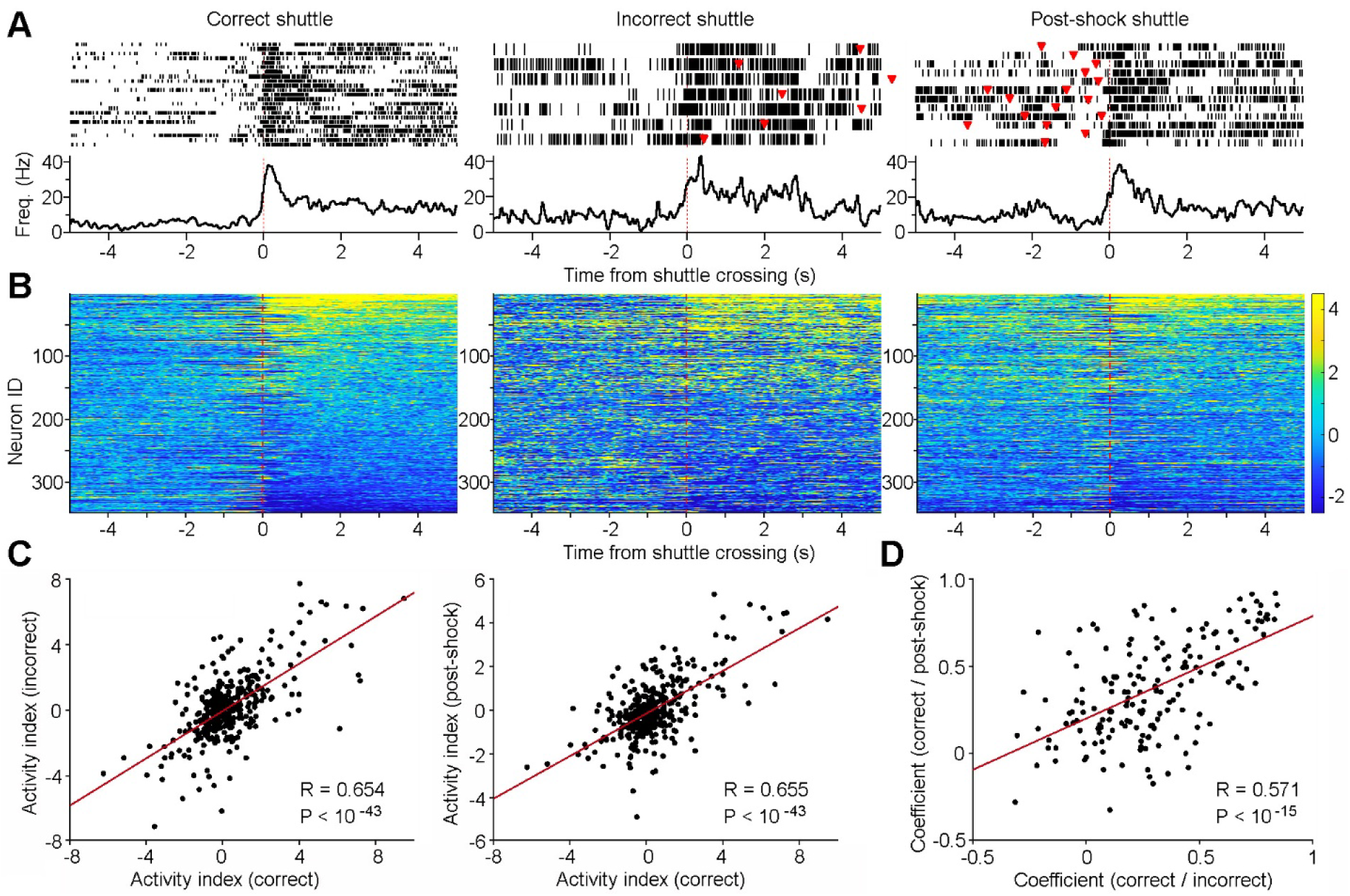
ACC neurons monitor actions independent of outcomes. **A,** Peri-event rasters (trials) and histograms of a representative ACC neuron during correct (left), incorrect (middle), and post-shock shuttles (right) within a session. Red triangles indicate shock administrations. Note that incorrect shuttles are followed by a second shuttle after animals receive footshocks, and approximately half of the post-shock shuttles are preceded by incorrect shuttles. **B,** Heatmaps showing the activity of individual ACC neurons (n = 348) during correct (left), incorrect (middle), and post-shock shuttles (right). Neurons are arranged in the same order across the three heatmaps. Color bar indicates z-scored activity. Note that the number of incorrect shuttles in a session is often ≤7, leading to greater variability in mean activity. Only sessions with ≥4 incorrect shuttles are included in the analysis. **C,** Activity indexes of individual ACC neurons between correct and incorrect shuttles (left), and between correct and post-shock shuttles (right). Activity index is defined as: Activity Index = Mean^post-shuttle^ – Mean^pre-shuttle^, where Mean^pre-shuttle^ and Mean^post-shuttle^ are the mean z scores calculated between −5–0 and 0–5 s, respectively, as shown in B. **D,** Correlation coefficients of the activity (−5–5 s) between correct and incorrect shuttles (x axis) and between correct and post-shock shuttles (y axis). Only the top and bottom quartiles of ACC neurons (as shown in B) are used for the analysis.

### ACC neurons monitor *action state* and *action content*

Since post-action ACC activity was outcome-independent, we next asked which other features of the task ACC activity might differentiate. Leveraging the two distinct shuttle responses within the task, we examined whether ACC neurons responded differently between rooms A→B shuttles (in response to sound A) and rooms B→A shuttles (in response to sound B). Our analyses identified two major groups of ACC neurons based on their responses to these shuttles. The first group showed indiscriminate response, exhibiting either increased or decreased activity without differentiating between A→B and B→A shuttles (Fig. 7: Neurons 1&2). We propose that these ACC neurons encode an *action state*, a variable representing the change of behavioral state in response to the auditory cues.

**Figure 7.**
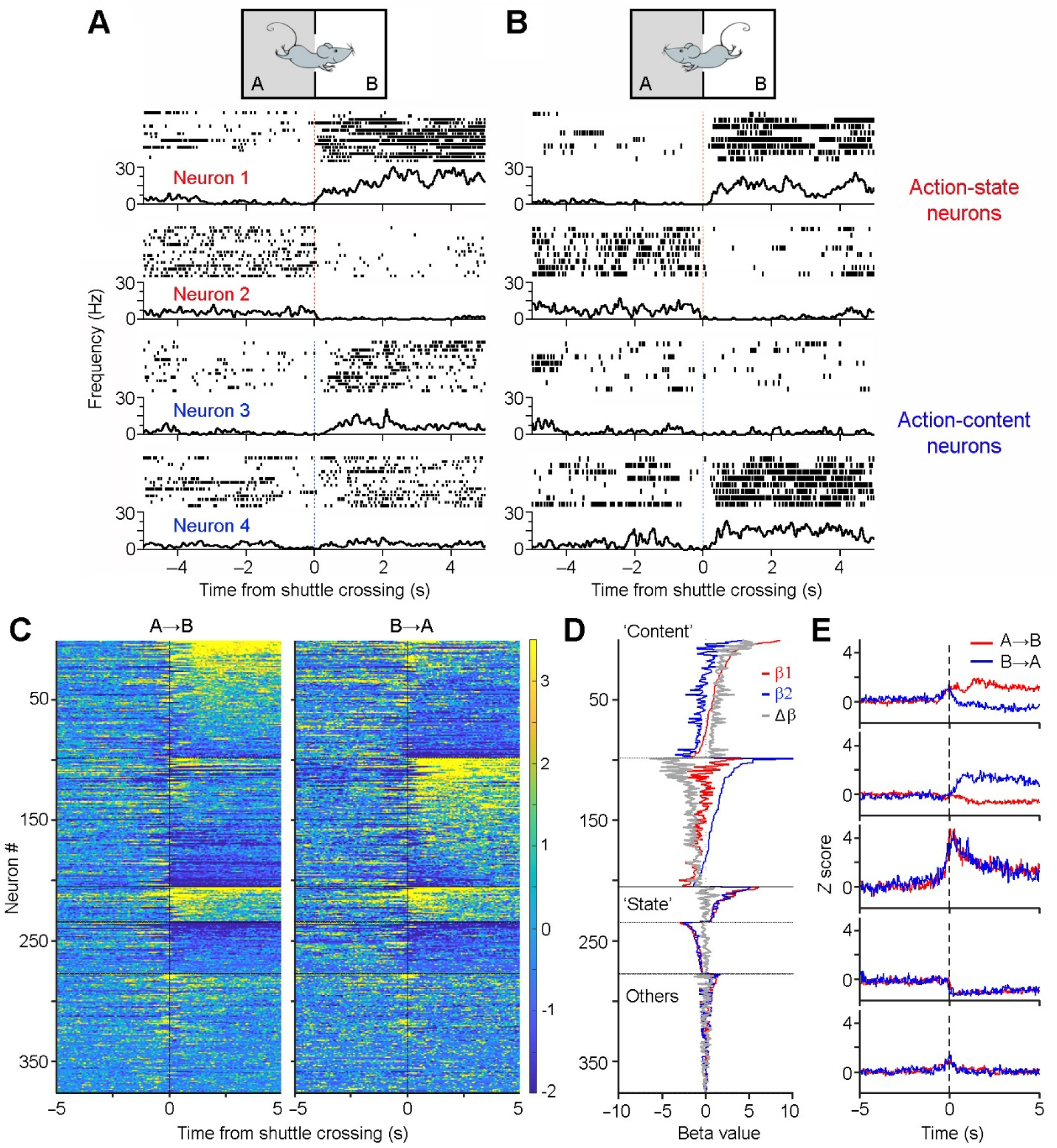
ACC neurons monitor *action state* and *action content*. **A&B,** Peri-event rasters (trials) & histograms of four simultaneously recorded ACC neurons during two sets of shuttles: rooms A→B shuttles (A) *vs.* rooms B→A shuttles (B). Notably, neurons 1&2 exhibit indiscriminate responses, either increasing or decreasing their activity after shuttles, thereby monitoring *action state* changes. In contrast, neurons 3&4 are selectively activated in one set of the shuttles, thereby monitoring *action content* (i.e., rooms A→B *vs.* B→A shuttles). **C**, Z-scored activity of all ACC neurons (n = 376) during A→B shuttles (left) and B→A shuttles (right). The color bar indicates z score. Neurons are categorized and sorted according to coefficient β or Δβ values (see Methods): Category 1 (n = 98), β₁ > β₂, sorted by β₁; Category 2 (n = 107), β₁ < β₂, sorted by β₂; Categories 3&4 (n = 29/43), both β₁ and β₂ are significantly positive or negative, sorted by the combined magnitude of β₁ and β₂. **D**, Coefficient values β₁, β₂, and Δβ (β₁–β₂) values are shown in red, blue, and gray, respectively, for individual ACC neurons ordered in the same sequence as shown in C. **E**, Mean activity for the five major neuronal categories (as discussed in C) during A→B (red) and B→A (blue) shuttles.

In contrast, the second group of ACC neurons exhibited discriminative responses between rooms A→B and B→A shuttles. These neurons either responded selectively in one set of shuttles or showed different levels of responses (activation or inhibition) between the two sets of shuttles (Fig. 7: Neurons 3&4; Fig. S6). We propose that these ACC neurons encode *action content*, a variable representing distinct action information (i.e., shuttling from rooms A→B *vs.* B→A).

To quantify *action-state* and *action-content* neurons, we performed a generalized linear model (GLM)-based analysis of ACC activity surrounding shuttle responses (see Methods). Based on coefficient β or Δβ value differences, most ACC neurons were classified as either *action-content* neurons (54.5%; Categories 1&2) or *action-state* neurons (19.1%; Categories 3&4; Fig. 7 C–E). These findings suggest a prominent role for post-action ACC activity in encoding rich information about preceding *action state* and *action content*.

Notably, ACC activity does not resemble place cell activity observed in the hippocampus [32], as evidenced by our analysis of spike activity from the intertrial-interval periods (Fig. S7). This aligns with prior findings that ACC neurons do not simply encode spatial information [33], although *action-content* ACC neurons likely incorporate spatial variables, given their shuttle-route response selectivity.

To determine if the post-action ACC neuronal population activity can decode shuttle contents (rooms A→B *vs.* B→A shuttles), we implemented a machine-learning approach. Specifically, we trained binary support vector machine (SVM) classifiers and performed cross-validations (Fig. 8A). Our results revealed that the post-shuttle population ACC activity was highly effective in decoding the animal’s preceding actions of either A→B *vs.* B→A shuttles, reaching a decoding accuracy close to 90% on average (Fig. 8B). Although the pre-shuttle ACC activity can also decode the shuttle content, it did so with much lower accuracy (Fig. 8B). Importantly, decoding accuracy was reliant on *action-content* neurons, as their removal significantly reduced decoding accuracy, whereas removal of non-*action-content* neurons had no effect (Fig 8C). These results persisted even when neuron category counts were matched across sessions in a pseudo-ensemble decoding analysis (Fig. S8). Only *action-content* neurons could reliably decode shuttle content, confirming a functional dissociation at the population level (Fig. S8). These findings provide compelling evidence supporting post-action ACC activity in encoding detailed information about the animal’s preceding actions.

**Figure 8.**
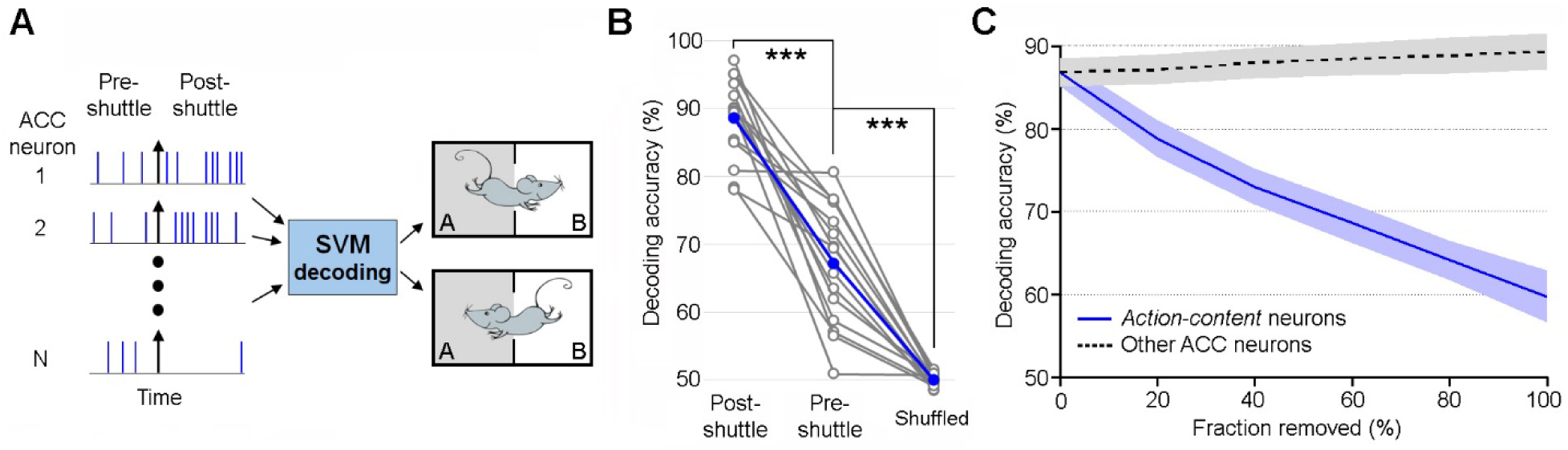
Post-shuttle ACC neuronal population activity decodes *action content*. **C,** Schematic diagram of support vector machine (SVM) decoding. ACC neuronal population activity from pre-shuttle period (−5–0), post-shuttle period (0–5), or shuffled spikes is used to train the decoder and subsequently distinguish between action content (rooms A→B *vs.* B→A shuttles). **B**, Mean decoding accuracy (blue line) and individual decoding accuracies for 15 sessions (grey lines). *Friedman test (*P < 0.001) and *post-hoc Wilcoxon signed-rank test with Bonferroni correction* (**\*\*\***P < 0.001). **C**, SVM decoding accuracy across all 15 sessions as a function of the fraction of neurons removed (20% per step), applied separately to *action-content* neurons (blue) or the remaining neurons (grey) within each session (P < 0.001 for each comparison between the two removals; *Wilcoxon signed-rank test*). Shaded areas denote ± s.e.m.

### Post-action ACC activity influences future performance

We next investigated whether post-action ACC activity following a trial influenced performance on the subsequent trial, which occurred ∼1 min later. Specifically, we divided correct-shuttle trials into two groups (Fig. 9A): those followed by correct trials (including both correct stays and shuttles), and those followed by incorrect trials (including incorrect stays and shuttles). Our results revealed that post-action ACC activity was notably higher when it preceded correct trials rather than incorrect ones (Fig. 9B). The strength of post-shuttle ACC activity may reflect task engagement, with greater engagement facilitating learning. Statistically, the top one third of the most responsive ACC neurons exhibit significantly higher activity that preceded correct trials than incorrect ones (Fig 9C). This correlation between higher ACC activity and future correct performance suggests that post-action ACC activity may contribute to trial-to-trial behavioral adjustment that underlies associative learning [34].

**Figure 9.**
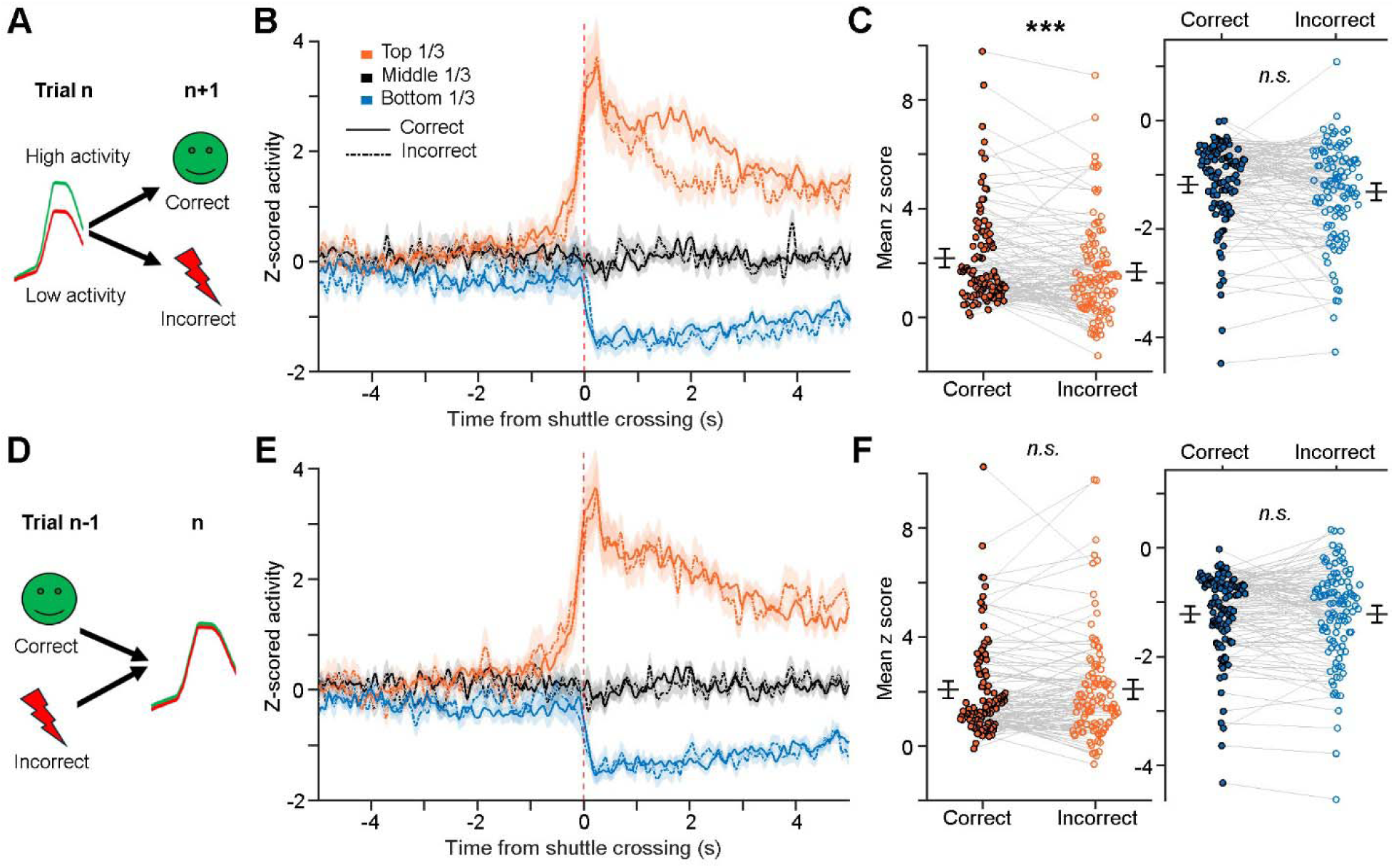
Post-action ACC activity influences future performance within a task session. **A,** Schematic illustration. **B**, Mean activity (± s.e.m.) of post-shuttle activated neurons (orange lines; top 1/3), inhibited neurons (blue lines; bottom 1/3), and remaining ACC neurons (black lines; middle 1/3), which preceded either correct trials (solid lines) or incorrect trials (dashed lines). **C,** Further comparison of the activation strength for post-shuttle activated neurons (left) and inhibited neurons (right) between the two conditions. Each pair of dots indicates an ACC neuron. Two-way ANOVA, Interaction: F_2, 328_ = 5.69; P = 0.004; Simple effect for Top1/3: ***P < 0.001; Effect Size: Cohen’s d = 0.33. **D–F,** Similar to A–C, except that the comparison is based on the status of the preceding trials. Mean z scores in C and F were calculated between 0–5 s after shuttle crossings. Two-way ANOVA revealed no-significant difference. n.s., non-significant.

As a control, we also examined whether the status of the preceding trial influenced post-action ACC activity in the current trial. Specifically, we divided correct-shuttle trials into two groups: those preceded by correct trials, and those preceded by incorrect ones (Fig. 9D). Our results showed no difference in post-action ACC activity between the two conditions (Fig. 9 E&F). This finding was expected, as both positive and negative reinforcement can similarly contribute to associative learning and neural plasticity.

## Discussion

Our newly designed discrimination–avoidance task is unique in that it allows us to disentangle the roles of ACC neurons across several proposed functions, including value encoding, decision-making, action monitoring, and outcome tracking. First, this task requires animals to discriminate both sensory cues and environmental contexts. Unlike established tasks that often assign fixed positive or negative values to cues, the cues in our task are not inherently associated with valence. Instead, their meaning is dynamically determined by the animal’s location (context) at the time of cue presentation. By removing valence from the cues, this design helps disentangle the ACC’s potential role in value encoding from other cognitive functions. Second, this task involves robust, ethologically relevant actions (i.e., shuttles), unlike many established paradigms that rely on less naturalistic behaviors such as saccades or lever presses. Finally, the clear temporal separation between actions and outcomes helps disentangle the ACC’s roles in action monitoring *vs.* outcome tracking.

Utilizing this task, we find that the ACC primarily encodes post-action variables. Specifically, ACC neurons exhibit robust post-shuttle responses across various conditions, including correct, incorrect, and post-shock shuttles. Despite conditions signaling distinct outcomes (presumed safety *vs.* uncertainty *vs.* safety), the response properties of the ACC neurons remain consistent. Moreover, very few ACC neurons respond directly to positive outcomes (safety) or negative outcomes (shocks). Together, these findings indicate that post-shuttle ACC activity primarily monitors animal’s most recent actions, rather than tracking the outcomes or values associated with those actions [21, 29, 30].

Our results further reveal two distinct groups of ACC neurons that encode different aspects of actions: *action state* and *action content*. Action state appears to represent the animal’s preceding choice of whether an action was taken, updating changes in behavioral state within the ongoing task. In contrast, *action content* represents the specific actions taken (i.e., shuttles from rooms A→B *vs.* B→A), preserving detailed action information. The response selectivity of *action-content* ACC neurons reinforces the notion that ACC activity monitors preceding actions rather than action-associated outcomes or values, which are uniform across these actions. Notably, our findings do not support an alternative interpretation that *action content* reflects correct *vs.* incorrect shuttles. Theoretically, categorizing actions solely as correct or incorrect may not be necessary, as both positive and negative outcomes can guide future actions and contribute to learning [30]. Potential neural networks mediating the sustained post-action ACC activity include thalamic and retrosplenial inputs, given the well-established roles of thalamocortical circuits in sustaining cortical activity [35, 36] and the retrosplenial cortex in processing spatial and directional information [37–40].

Our study also reveals that ACC neurons play a limited role in encoding pre-action variables associated with decision-making or planning, as evidenced by their minimal responses to auditory cues and the modest activity changes prior to shuttle initiation. These findings align with recent research, showing that ACC neurons are mainly involved in post-decisional information processing rather than decision-making itself [18–20]. Nevertheless, substantial evidence from other studies supports the ACC’s involvement in value coding or value updating, as reflected in its differential responses to discriminative sensory cues that precede decisions [25, 31, 41]. One possible explanation for the discrepancy in our findings is that the sensory cues used in our task are not intrinsically linked to specific values. Although these cues play a crucial role in driving go (shuttle)/no-go (stay) decisions, their values are dynamic and vary across trials. This lack of value association may explain why we observe minimal response of ACC neurons to these cues. In contrast, the ACC activity reported in previous studies likely reflects the stable value of the cues [31, 42, 43]. It remains to be determined if distinct ACC subpopulations may be responsible for value encoding *vs.* action monitoring, or if the same ACC neurons multiplex across these functions.

Our results suggest that ACC activity is not directly associated with locomotion. First, both *action-state* and *action-content* neurons tend to show sustained activity even when the animals remain immobile after completing shuttle behaviors, suggesting that their activity is not driven by locomotion. Furthermore, *action-content* neurons are selectively engaged in only one of the two shuttle categories, either rooms A→B or B→A shuttles. Therefore, differences in neuronal activity are unlikely to reflect locomotor differences, given that both shuttle types involve similar movement patterns. Finally, we show that only a small fraction of neurons (14.6%) exhibits locomotion speed-correlated activity. Overall, these results suggest that post-action ACC activity reflects information about preceding actions, independent of the animal’s current movement.

One caveat of our study is that the discrimination–avoidance task requires weeks of training in mice. By the time they master the task, ACC activity may reflect modified neural circuits. Investigating ACC activity during early phase of learning, such as by introducing a new pair of cues or contexts, could provide further insights into ACC’s role in learning and cognitive processes. Additionally, previous studies have highlighted ACC’s key role in reversal learning [44–47]. Future research examining how ACC neurons respond when task rules are reversed could provide further insight into this function. Histological verification of the recording sites revealed that a small subset of recordings resided at the border between the ACC and motor cortex (Fig. S2). Although most recordings were situated within the ACC, it remains possible that some of the neural responses described originated from motor cortex neurons and may influence these findings. Finally, a limitation of the current study is the lack of evidence for the causal role of post-action ACC activity in complex associative learning. Future investigations using closed-loop strategies to selectively disrupt ACC activity during the post-action phase could help address this question.

Lastly, our findings suggest that post-action ACC activity plays a key role in shaping future behavior. Specifically, we find that increased post-action ACC activity is linked to future performance in subsequent trials, highlighting the ACC’s role in facilitating learning and guiding behavior. The level of post-shuttle ACC activity may reflect task engagement, with greater engagement facilitating learning and improving future performance. This aligns with previous research demonstrating the ACC’s critical role in complex and flexible learning, including discriminative avoidance learning [48], discriminative extinction learning [49], reversal learning [44–47], task switching [25], trial-to-trial behavioral adaptation [34], and action–outcome associative learning [28–30]. We speculate that both *action-state* and *action-content* ACC neurons contribute to complex associative learning through extended signaling of action-relevant information, thereby bridging cue, action, and outcome associations (Fig. 10). Specifically, *action-state* ACC neurons signal whether an action was taken, while *action-content* ACC neurons provide specific details about what the action was—the content. This action-related information, eventually, is integrated with cue- and outcome-related variables to form complex cue–action–outcome associations that guide future behavior (Fig. 10). Without the ACC, action-relevant information could be lost after action execution, thereby disrupting cue–action–outcome associative learning. Given our key finding that the ACC primarily encodes action-related variables rather than cue- or outcome-related information, we speculate that the integration of these complex associations takes place in other higher cognitive areas, such as the prelimbic cortex and other medial prefrontal subregions [50–52].

**Figure 10.**
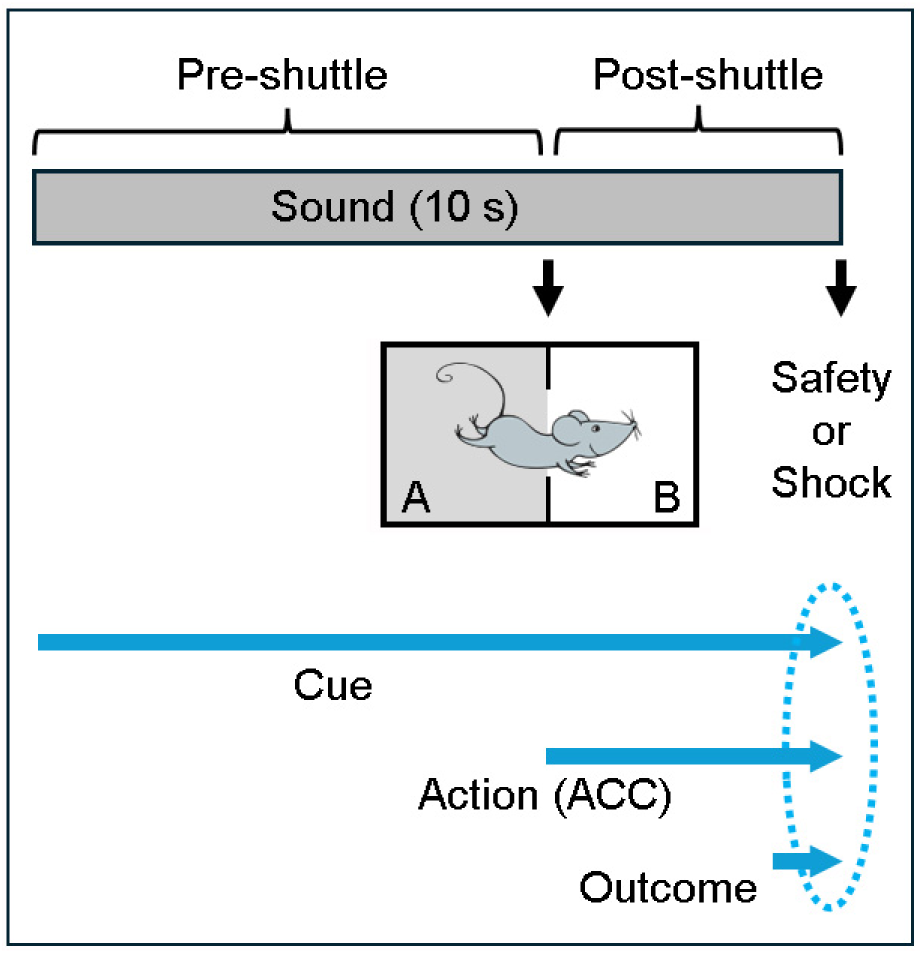
Proposed model of cue–action–outcome associative learning. **Top:** Four distinct phases of information coding: pre-shuttle, shuttle, post-shuttle, and outcome phases. **Bottom:** Integration of cue, action, and outcome information occurs when all three components are concurrently available (indicated by the oval). Notably, without the ACC, action-relevant information could be lost after action execution, thereby disrupting cue–action–outcome associative learning.

## Methods

### Mice

Male C57BL/6 mice (Jackson Laboratory, stock #000664) were used in this study. The mice were 8–10 weeks old at the start of discrimination–avoidance task training. All mice were group-housed (2–4 mice per cage; 40 × 20 × 25 cm) with corn cob bedding and cotton nesting material, except that after electrode implantation surgery, mice were singly housed. They were maintained on a 12-h light/dark cycle with *ad libitum* access to food and water. All experimental procedures were approved by the Institutional Animal Care and Use Committees at Drexel University and adhered to the National Research Council’s Guide for the Care and Use of Laboratory Animals.

### Sounds and shuttle box

Two 10-s auditory cues, 5-kHz tone at ∼75 dB and white noise at ∼65 dB, were chosen for sound discrimination. Both sounds included 50-ms shaped rise and fall times to reduce abruptness and minimize potential startle responses. The shuttle box used in the experiment was a square chamber measuring 25 × 25 × 32 cm, with a 36-bar shock grid floor, illuminated by lights inside sound-attenuating cubicles (64 × 75 × 36 cm) equipped with speakers (*Med Associates*). The shuttle box was divided at the midline by a plastic divider into two rooms. These two rooms were slightly modified for discrimination purposes: one had two black walls, one white wall, and one transparent wall, while the other room had two white walls, one metal wall, and one transparent wall (Fig. S1). The divider had a 2-inch opening in the center to allow the mice to move freely between the rooms. Animals’ behaviors were recorded using Video Freeze software (*Med Associates*) [53].

### Discrimination–avoidance task

Prior to training, all mice underwent two daily handling sessions (∼10 min each). Once training began, the mice received one training session per day, 5 days per week. In the task, the mice were trained to discriminate between two distinct sounds (A and B) and shuttle between two adjacent rooms (A and B) within a shuttle box to avoid potential footshocks. Specifically, sounds A and B signaled electric shocks in rooms A and B, respectively (Fig. 1A). During training, the mice were first allowed to freely explore the shuttle box for 2 min before trials began, except on the first day of training, where the free exploration period was extended to 10 min.

The training procedure consisted of two phases: pre-training and training. Pre-training phase: mice underwent five daily sessions of sound alternation trials 5/5 (AAAAABBBBB …). Mice that did not exhibit shuttling responses during the first session were excluded from further training. Training phase: mice underwent 10 daily sessions (5 sessions per week) of alternation trials in a pseudorandom order (ABBABAAB …).

Each training session comprised 50 trials, 60 s apart. On incorrect trials, up to 10 scrambled electric shocks (0.5 mA; 0.1 s) were administered starting at sound terminations and continued for an additional 18 s (1 shock every 2 s), except during the pre-training phase, where up to 20 scrambled electric shocks were administered. Shocks were terminated once the mice navigated to the adjacent safe room. The mouse’s success rate in avoiding shocks at sound terminations was defined as: Success rate (%) = (Correct stays + Correct shuttles) / All trials.

### Real-time location detection

We employed MATLAB functions to detect animal location and subsequently control shock administration (Fig. S1). Specifically, *Med Associates* Video Freeze software recorded real-time video footage of the animal’s behaviors [53]. During each trial, MATLAB captured screenshots of the ongoing video at sound terminations and every 2 s thereafter during the shock period. MATLAB then performed background extraction on the captured images to determine which room the mouse was located in. To deliver a shock, MATLAB sent a “shock” signal to an *Arduino UNO* circuit board, which relayed that signal to *Med Associates*, triggering shock administration.

### Stereotaxic surgery

Mice that had completed 10 training sessions and surpassed a success rate of 70% during discrimination–avoidance tasks were used for surgery. In brief, mice were anesthetized with ketamine/xylazine mixture (∼100/10 mg/kg, i.p.) and maintained on a heating pad at 37°C. Following that, mice received an intra-ACC implantation of a custom-made electrode array (8 tetrodes) [54, 55], and the implant was secured to the skull with stainless screws and resin ionomer (*DenMat*). The coordinates used were AP 1.0 mm, ML 0.4 mm, and DV 1.0 mm.

### *In vivo* recording during discrimination–avoidance tasks

We used tetrodes for recording [54, 55]. Each tetrode consisted of four wires (90% platinum 10% iridium; ∼18 μm diameter; *California Fine Wire*). Neural signals were preamplified, digitized, and recorded using a *Blackrock Neurotech* CerePlex, while the animals’ behaviors were simultaneously recorded. Spikes were digitized at 30 kHz and filtered between 600–6000 Hz. The recorded spikes were manually sorted using *Plexon* Offline Sorter [56, 57], with key datasets verified by a second experimenter. Tetrode arrays were gradually lowered through a microdrive [58, 59] to record at multiple depths within the ACC, with 2–4 depths from each animal (DV ∼1.0–1.3) used for analysis.

Overall, ACC spikes from five mice across 15 sessions were analyzed, with neuron counts of 20/23/29/32 (mouse #1; 4 sessions), 17/20/22 (mouse #2; 3 sessions), 28/24 (mouse #3; 2 sessions), 33/25/21/27 (mouse #4; 4 sessions), and 22/25 (mouse #5; 2 sessions), respectively. The total number of correct/incorrect shuttles used for analysis are 19/5, 19/4, 21/5, 20/4 (mouse #1); 20/7, 23/7, 20/7 (mouse #2); 19/4, 16/2 (mouse #3); 26/4, 23/4, 17/6, 25/5 (mouse #4); 20/5, and 17/4 (mouse #5), respectively. For Figs. 2–5 & 7–8, all recorded neurons (n = 376) from the 15 sessions were included in the analyses. For Fig. 6, however, one or two sessions were excluded because they contained too few trials of certain types (n ≤ 3).

The discrimination–avoidance task was similar to that during the training phase, except that five direct-current electric shocks, which minimize electromagnetic artifacts, were administered starting at sound terminations and continued for 8 s. This shorter shock period allowed the mice more time to move freely within the shuttle box during the 42-s inter-trial intervals, when no consequences were administered. Additionally, each task session included a 5-min free exploration period before trials began. In some sessions, a small number of trials were excluded from analysis, partly due to the electrode implant or recording cable occasionally interfering with the animal’s shuttle responses.

### Shuttle behavior analysis

We used DeepLabCut [60] to analyze animals’ locations during discrimination–avoidance tasks, with all locations determined based on body center positions. Shuttle crossing was defined as the body center crossing the midline opening of the shuttle box. Shuttle initiations and terminations were defined as the time points when the animal’s movement velocity deviated 1 s.d. above mean (Fig. 4A).

### Dimension reduction

We used Principal Component Analysis (PCA) to analyze the major activity patterns of ACC neuronal populations. The first three principal components (PCs) were used for 3-D visualizations. Specifically, the activity of each ACC neuron was first averaged during shuttle responses (bin size, 0.1 s) and z-scored. The z-scored activity of ACC neurons was then processed with PCA (*pca*, MATLAB).

### Event-locked modulation analysis

Neural activity was aligned to shuttle initiations, crossings, or terminations following standard peri-event procedures. For each neuron, firing-rate modulation (ΔFR) was computed as the difference between post-event (0–W ms) and pre-event (−W–0 ms) windows (W = 250–2000 ms), similar to prior event-related analyses [61]. For each event, significance was assessed using an empirical sliding-window null distribution generated within a baseline period (−20 to −10 s). Two-sided empirical p-values were computed, and neurons were considered significantly modulated if p < 0.01 and effect size |Δ| ≥ 0.6. Neurons were classified as event-specific, crossing-only, both, or non-significant based on the overlap of significant modulation. Differences in modulation prevalence were evaluated using exact McNemar tests [62].

### Speed tuning analysis and speed cell classification

Speed tuning was assessed using neural and behavioral data collected during a ∼5-min period of free movement in the shuttle box prior to task onset. During this period, animals freely explored the environment and exhibited a broad range of instantaneous running speeds. Running speed was extracted from behavioral tracking data and sampled at 50 Hz. Spike trains from individual neurons were binned at 20 ms resolution and converted to firing rates. Firing-rate time series were smoothed using a Gaussian kernel with a 250 ms window. To reduce potential confounds associated with immobility-related network states, time points with running speed below 2 cm/s were excluded from the analysis, following established procedures [63]. For each neuron, a speed score was defined as the Pearson correlation coefficient between the smoothed firing rate and the valid portion of the instantaneous running speed.

Statistical significance was assessed using a circular time-shift shuffle procedure. For each neuron, spike times were circularly shifted by a random offset greater than 30 s, preserving the temporal structure of firing while disrupting its alignment with behavior. The firing rate–speed correlation was recomputed for each shuffle, and this procedure was repeated 100 times to generate a null distribution of speed scores. Neurons were classified as speed-modulated if their observed speed score exceeded the 99th percentile or fell below the 1st percentile of the shuffle distribution (two-sided criterion). All analysis procedures were adapted from previous studies of speed-modulated neurons [63].

### PCA-based classification of ACC neuronal types

We used PCA to classify major types of ACC neurons based on their activity in reference to shuttle initiation. Specifically, the activity of each ACC neuron was first averaged surrounding shuttle responses (−5–5 s; bin size, 25 ms) and z-scored. The first three PCs were used in a hierarchical clustering algorithm (Linkage) to find the similarity (Euclidean distance) between all pairs of activity patterns in PC space, iteratively grouping the activity patterns into larger and larger clusters based on their similarity. Lastly, we set a distance-criterion to extract major clusters from the hierarchical tree [64].

### GLM-based classification of ‘shuttle-content’ and ‘shuttle-state’ neurons

For each neuron, we computed peri-event firing rates using 25-ms bins and normalized activity relative to a baseline window from −30 to −20 s preceding shuttle onset. Post-shuttle activity (0–5 s) was analyzed using a generalized linear model (GLM) with separate regressors for each event. Specifically, for Event1 (shuttle A→B) and Event2 (shuttle B→A) trials, we fit the model

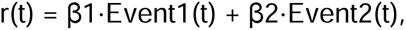

using a normal distribution with an identity link (*glmfit*, constant = ‘off’, MATLAB). We estimated coefficients β1 and β2 (corresponding to shuttle A→B and shuttle B→A, respectively) and assessed their significance using Wald tests against zero. To quantify event selectivity, we tested the contrast: Δβ = β1–β2. Resulting p-values were corrected for multiple comparisons using Bonferroni correction across regressors and neurons (α_bonf = 0.01/(3N), where N is the number of neurons in a session). Neurons were classified as *action-content* neurons if the corrected p-value for Δβ was significant and the absolute effect size exceeded a predefined threshold (| Δβ | > 0.5). Neurons were classified as *action-state* neurons if Δβ was not significant but both β1 and β2 were individually significant after correction. All analysis procedures were adapted from previous studies [65–67].

### Machine learning decoding

We used Support Vector Machine (SVM) classifiers to train ACC neuronal population activity to decode shuttle content (rooms A→B *vs.* B→A shuttles) and subsequently performed 10-fold cross-validations. Specifically, the total number of spikes from each ACC neuron, calculated during either the pre-shuttle (−5–0 s) or post-shuttle period (0–5 s), were used for training and testing the SVM classifier. For each dataset, we performed SVM classification training and cross-validation 100 times (*fitcsvm* and *crossval*, MATLAB), assigning the mean correct classification rate as the decoding accuracy. Notably, using shorter or longer time windows surrounding shuttle responses, such as 2.5 or 10 s, yielded similar conclusions (Fig. S9). Additionally, ACC neuronal population activity moderately decoded Room-A *vs.* Room-B stays (Fig. S9; SVM training and cross-validation repeated 500 times).

To assess the contribution of *action-content* neurons to decoding performance, neurons were progressively removed in 20% increments (Fig. 8C). Within each session, *action-content* neurons were ranked by | Δβ | in descending order, and the top fraction was removed at each step prior to re-estimating decoding accuracy. As a control, the same number of neurons was randomly removed from the Δβ non-significant pool. At each removal fraction, decoding accuracies obtained after *action-content* neuron removal were compared to control accuracies across sessions using a paired, two-sided, Wilcoxon signed-rank test.

### Pseudo-ensemble decoding of action identity

To quantify population-level encoding of *action contents*, we performed pseudo-ensemble decoding using trial-level firing rates. For each neuron, firing rate was computed separately for each trial within the post-shuttle window (0–5 s). For a given ensemble size (N neurons), pseudo-population vectors were constructed by randomly sampling one trial per neuron and concatenating their firing rates into an N-dimensional vector. Because neurons were recorded across different sessions, pseudo-ensembles were generated by independently sampling trials across neurons at each resampling iteration. Within each neuron, trial numbers were balanced across conditions by subsampling the larger condition to match the smaller (at least 4 trials). Trials were partitioned into training and test pools using a fixed holdout fraction (25%). Pseudo-population vectors were generated separately from the training and test pools to ensure independence between model fitting and evaluation. The SVM classifier was trained to discriminate *action contents* (A→B *vs.* B→A) using training population vectors. Decoding accuracy was computed exclusively on held-out test vectors (N = 10, 20, 30, 40, or 50). This procedure was repeated across multiple ensemble sizes and resampling iterations to obtain stable estimates of decoding performance. Shuffle controls were implemented by randomly permuting training labels while keeping test labels unchanged, thereby preserving population statistics while eliminating condition identity [62, 68].

### Spatial information analysis (Fig. S7)

To quantify spatial information coding, firing rate maps for individual neurons were constructed using 1 × 1 cm spatial bins in NeuroExplorer and smoothed with a Gaussian filter (filter width, 3 bins) before being exported to MATLAB for further analyses. Spatial information for each neuron was calculated using the following formula:

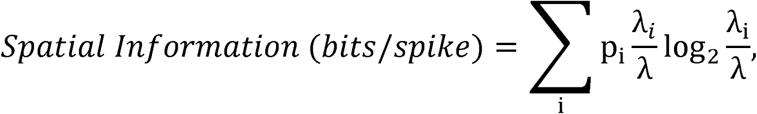

where λ_i_ is the mean firing rate in the i-th bin, λ is the overall mean firing rate, and p_i_ is the probability of the animal’s being in the i-th bin, as described previously [55, 69].

### Histology

To mark the final recording sites, we made electrical lesions by passing 10-s, 10-μA currents through multiple tetrodes. Mice were deeply anesthetized and intracardially perfused with ice-cold PBS or saline, followed by 10% formalin. The brains were removed and postfixed in formalin for at least 24 hours. The brains were sliced into coronal sections of 50-μm thickness using *Leica* vibratome. Brain sections were mounted with Mowiol mounting medium for microscopic examination of electrode array placements.

### Statistics

Sample sizes were based on previous similar studies [55, 64]. All statistics were conducted in SPSS 30.0. Statistical analyses include repeated measures ANOVA, the nonparametric Friedman test by post-hoc test (pair-wise Wilcoxon signed-rank test and Bonferroni correction), the Wilcoxon signed-rank test (paired), and Student’s *t* test (paired). All statistical tests are two-sided; P-values of 0.05 or lower were considered significant.

## Acknowledgements

This work was supported by the National Institutes of Health grants R01MH119102 (D.V.W.), R21MH134016 (D.V.W.), and F31MH134582 (A.F.H.).

---

**Supplementary Video 1. A representative correct shuttle response.**

**Supplementary Video 2. A representative incorrect shuttle response.**

**Fig. S1.**
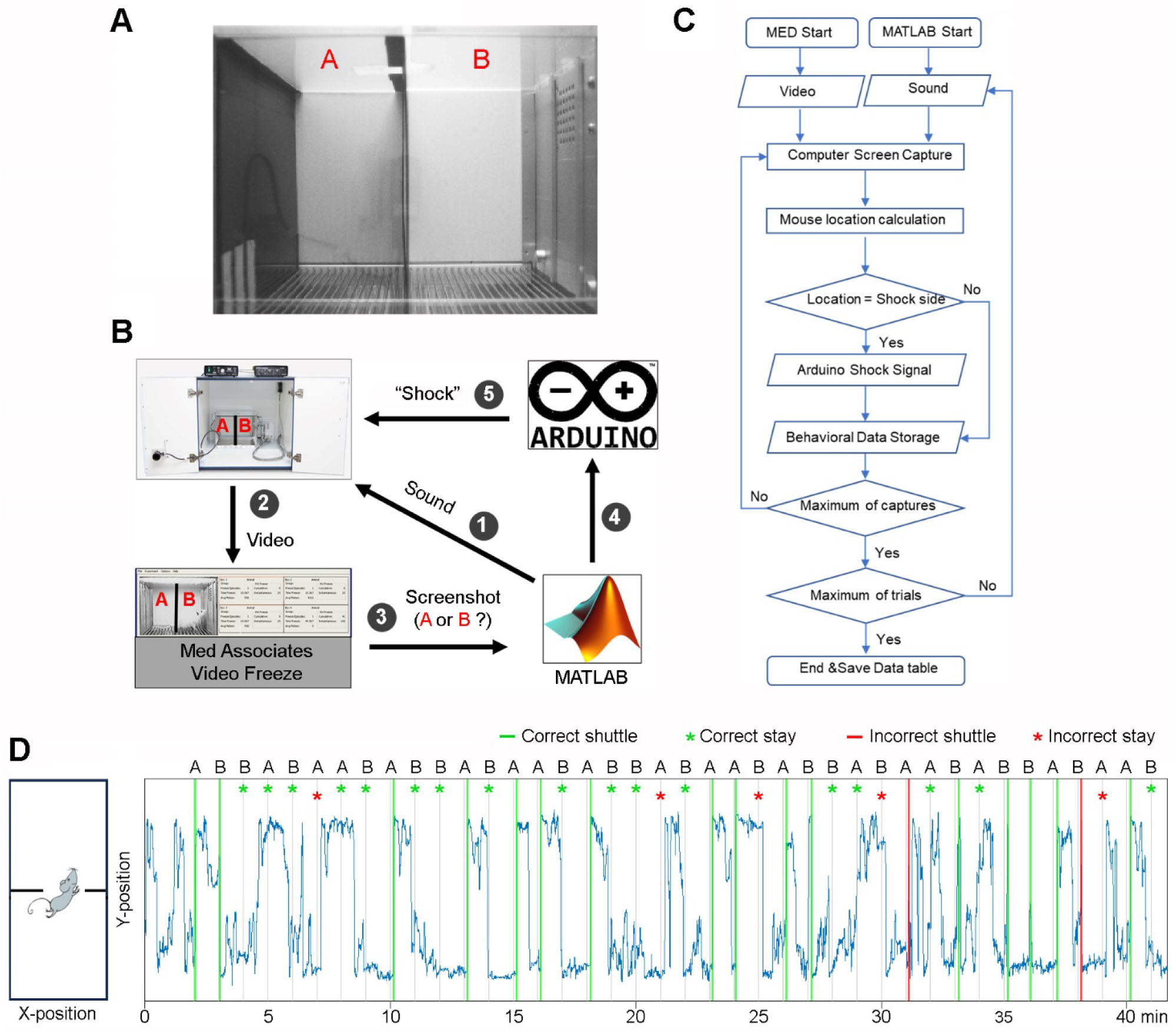
Experimental setup. **A,** The shuttle box used for behavioral training. Room A is configured with two black walls (left and right), one white wall (back), and a transparent front wall for video recording purposes. Room B is configured with two white walls (left and back), one metal wall (right), and a transparent front wall. **B,** Schematic diagram of the control setup utilizing MATLAB functions, with numbers indicating the sequence of control flow. **C,** A comprehensive flowchart illustrating the control setup as shown in B. **D,** Left, sounds A and B signal shocks in the bottom and top rooms of the shuttle box, respectively. Right, the Y position of a well-trained mouse in a ∼40-min session.

**Fig. S2.**
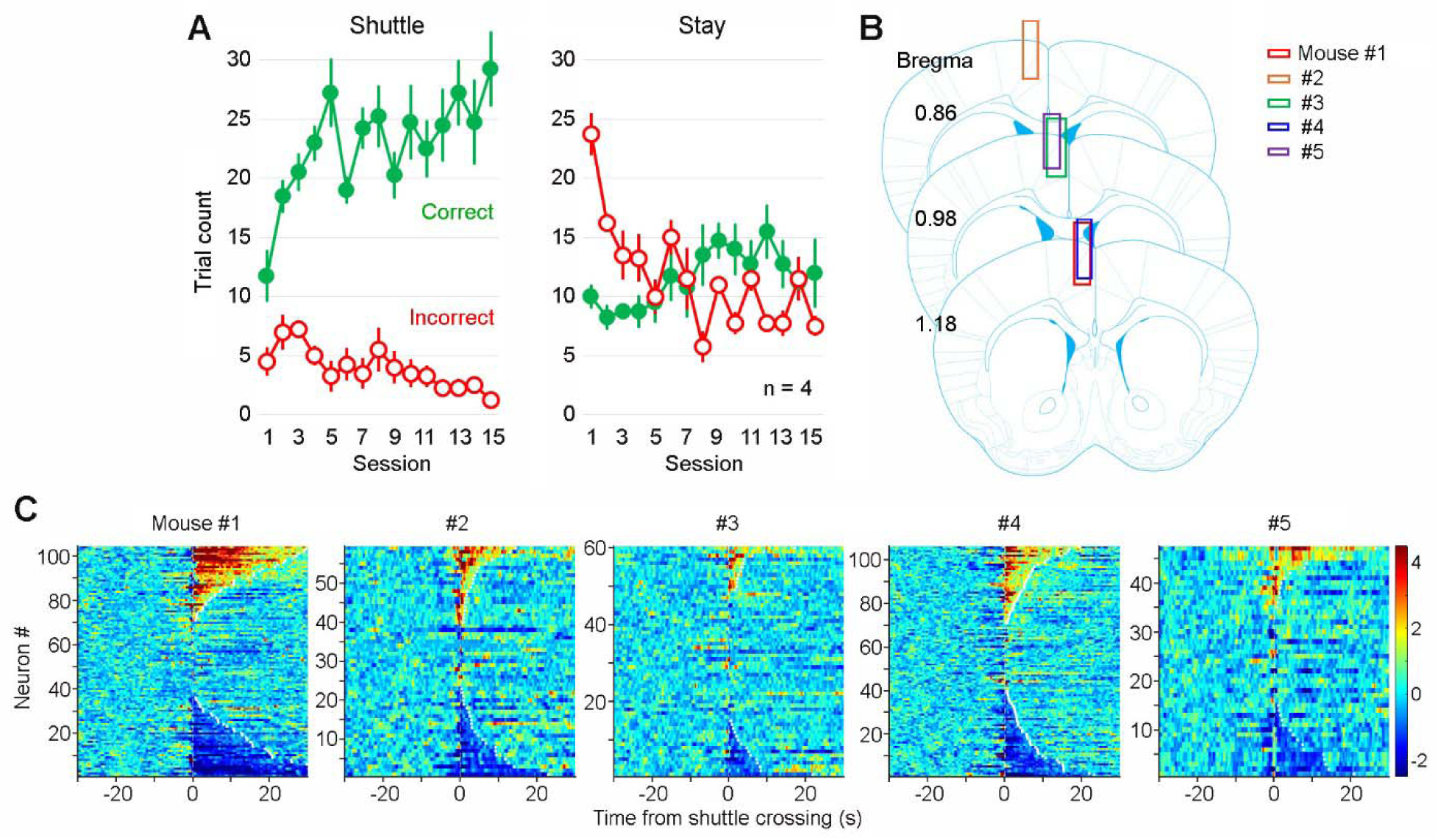
Extended training recording sites. **A,** Correct and incorrect trial counts for shuttle *vs.* stay trials across 15 training sessions. In shuttle trials, mice must move to the adjacent room before the sound ends to avoid shocks, whereas in stay trials, mice must remain in their current room to avoid shocks. **B,** reconstructed recording sites for individual mice. **C,** Heatmaps showing the activity of all recorded ACC neurons during correct-shuttle trials across individual mice.

**Fig. S3.**
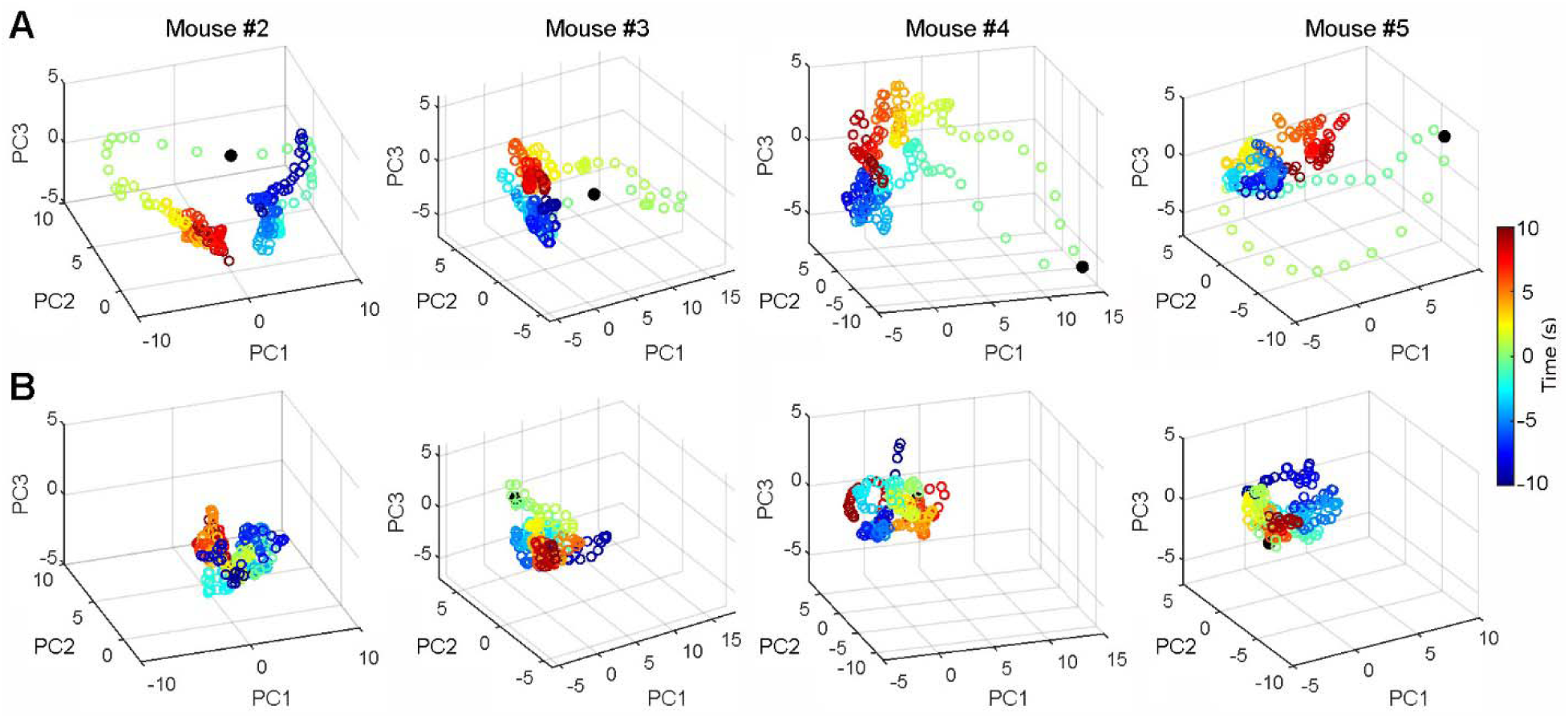
ACC neurons primarily respond during “shuttle” but not “stay” trials. **A&B,** Dimension reduction analysis (i.e., PCA) indicates robust changes in ACC neuronal population activity during correct shuttles (A) but not correct stays (B) from four representative recording sessions. Time “0” indicates shuttle crossings (A) or sound onsets (B), respectively. For more details, see Fig. 3B.

**Fig. S4.**
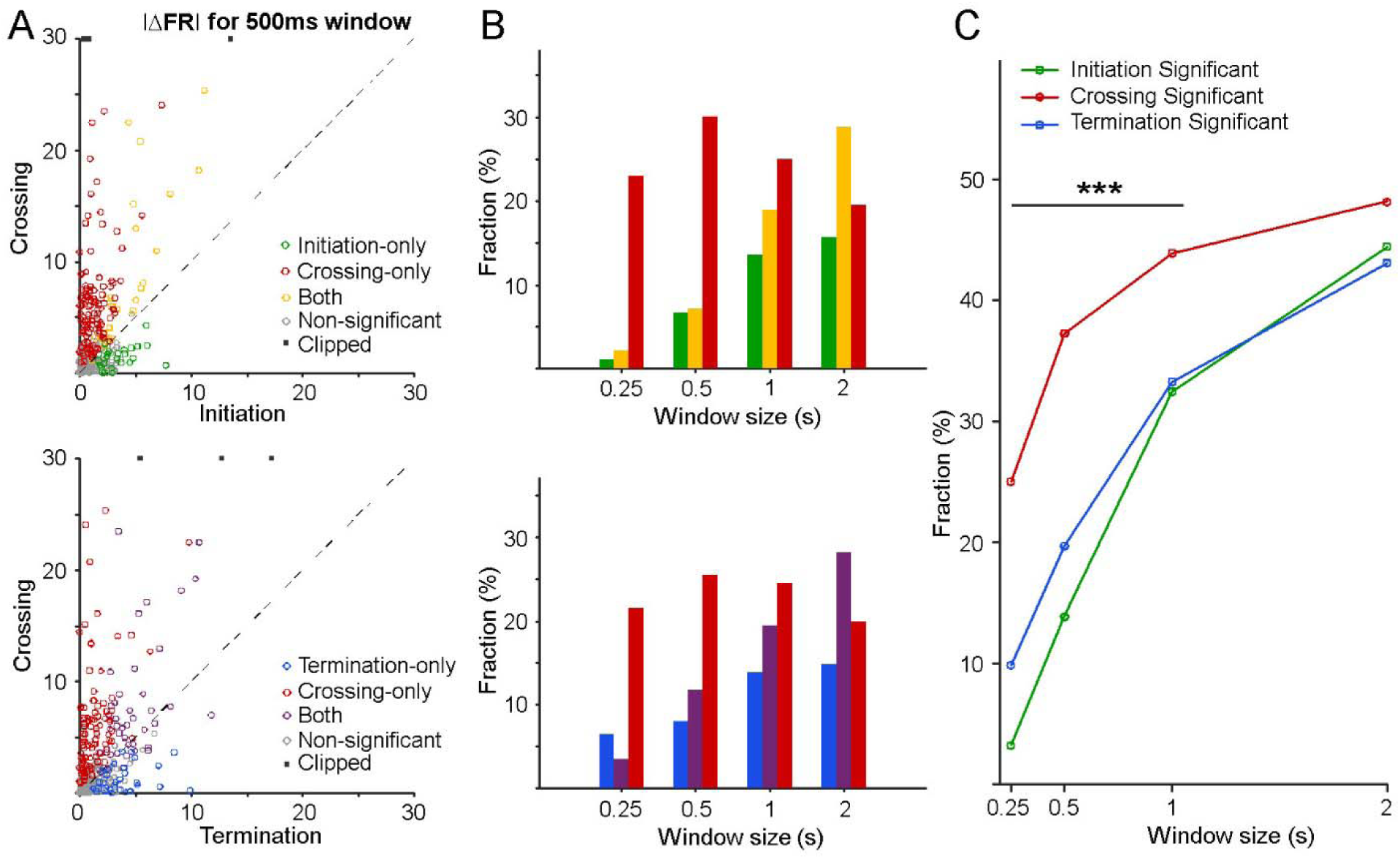
Event-locked modulation across shuttle initiations, crossings, and terminations. **A,** Scatter plots showing modulation magnitude (|ΔFR|) between pre- and post-event windows (500 ms; see Methods). Each dot represents one neuron. Top: initiation *vs.* crossing. Bottom: termination *vs.* crossing. Neurons were classified as event-specific (initiation-only, crossing-only, or termination-only), both, or non-significant based on empirical null testing. Points exceeding the plotting range are shown as dark squares. **B,** Fraction of neurons classified as event-specific, both, or non-significant across different time windows (0.25–2 s). **C,** Fraction of neurons significantly modulated by initiation, crossing, or termination across different time windows. Significance was assessed using an empirical sliding-window null distribution; stars indicate McNemar test results comparing initiation- *vs.* crossing-, and termination- *vs.* crossing-modulated fractions (***p < 0.001). Note that as the analysis window was expanded, differences between event types became less distinct, and overlap between them increased, likely reflecting the rapid execution of the shuttle behavior (typically completed within ∼2 s), which temporally compresses event-related neural dynamics.

**Fig. S5.**
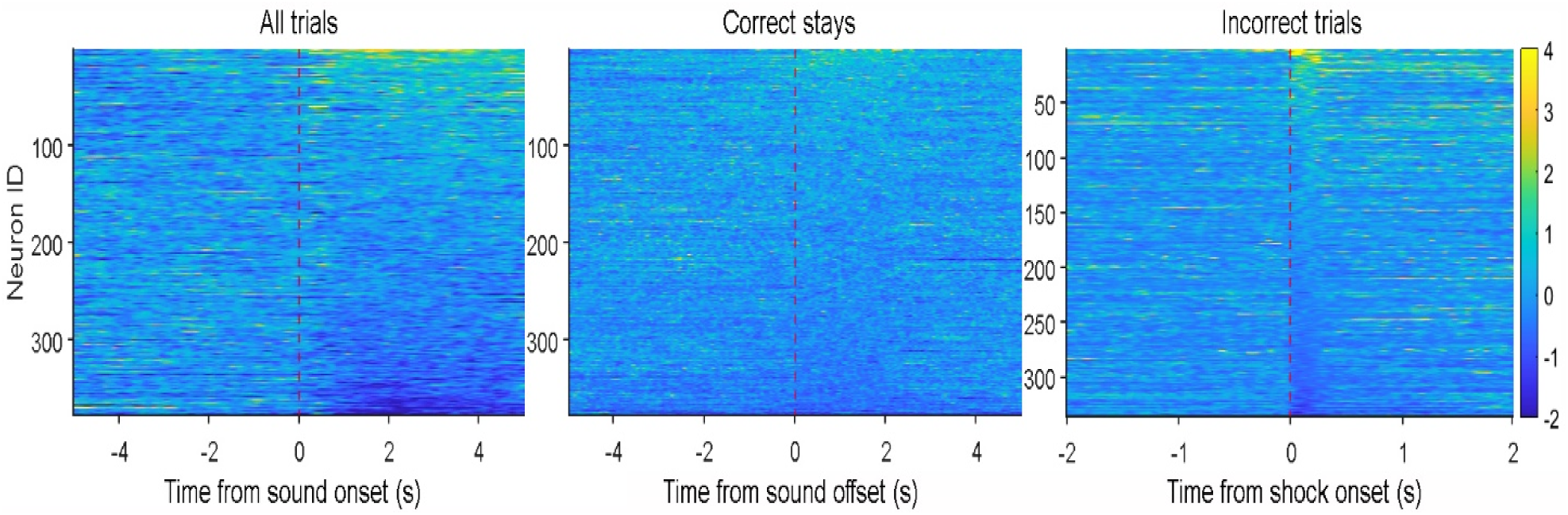
ACC shows limited response to cue, stay trials, and footshocks. Left, Heatmap showing the activity of individual ACC neurons (n = 348) in relation to auditory cue onset. Middle, Heatmap showing the activity of individual ACC neurons (n = 348) during stay trials. Right, Heatmap showing the activity of individual ACC neurons (n = 336) during footshock. Note, footshock trials with a shuttle response within 1 s shock onset were excluded to avoid shuttle response confound. As a result, some behavioral sessions were excluded due to insufficient trial numbers.

**Fig. S6.**
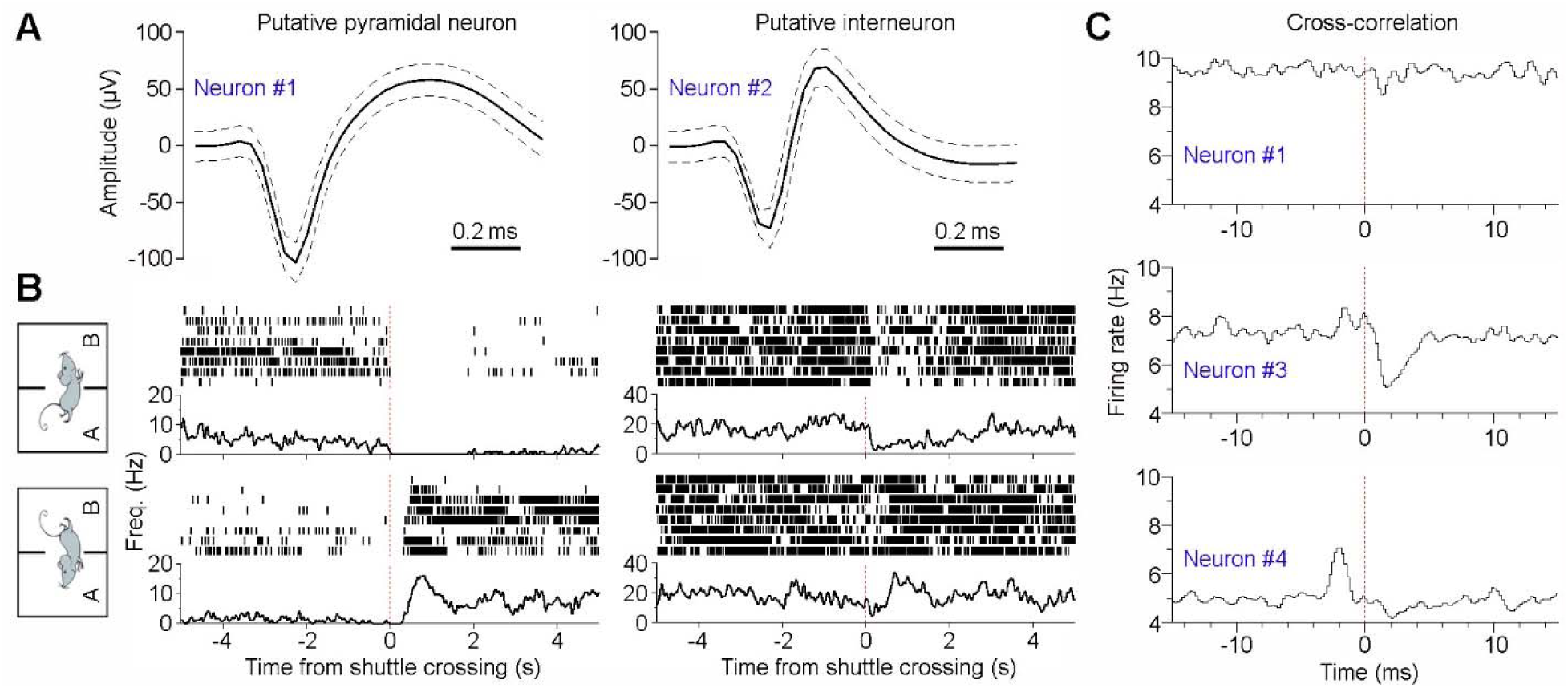
ACC pyramidal neurons and interneurons both monitor *action content*. **A,** Spike waveform (mean ± s.d.) of two representative ACC neurons: one putative pyramidal neuron and one interneuron. **B,** Peri-event rasters (trials) & histograms of the same two ACC neurons surrounding shuttle responses. Both neurons exhibit differential activity changes that discriminate between rooms A→B (top panels) *vs.* B→A shuttles (bottom panels). **C,** Cross-correlation histograms between the putative interneuron (Neuron #2; the same as shown in A) and three other pyramidal neurons (Neurons #1, #3, and #4). Neuron #4 appears to excite Neuron #2, which in turn inhibits Neuron #3, as indicated by short-latency (∼2 ms) excitatory or inhibitory interactions. The four ACC neurons were recorded simultaneously.

**Fig. S7.**
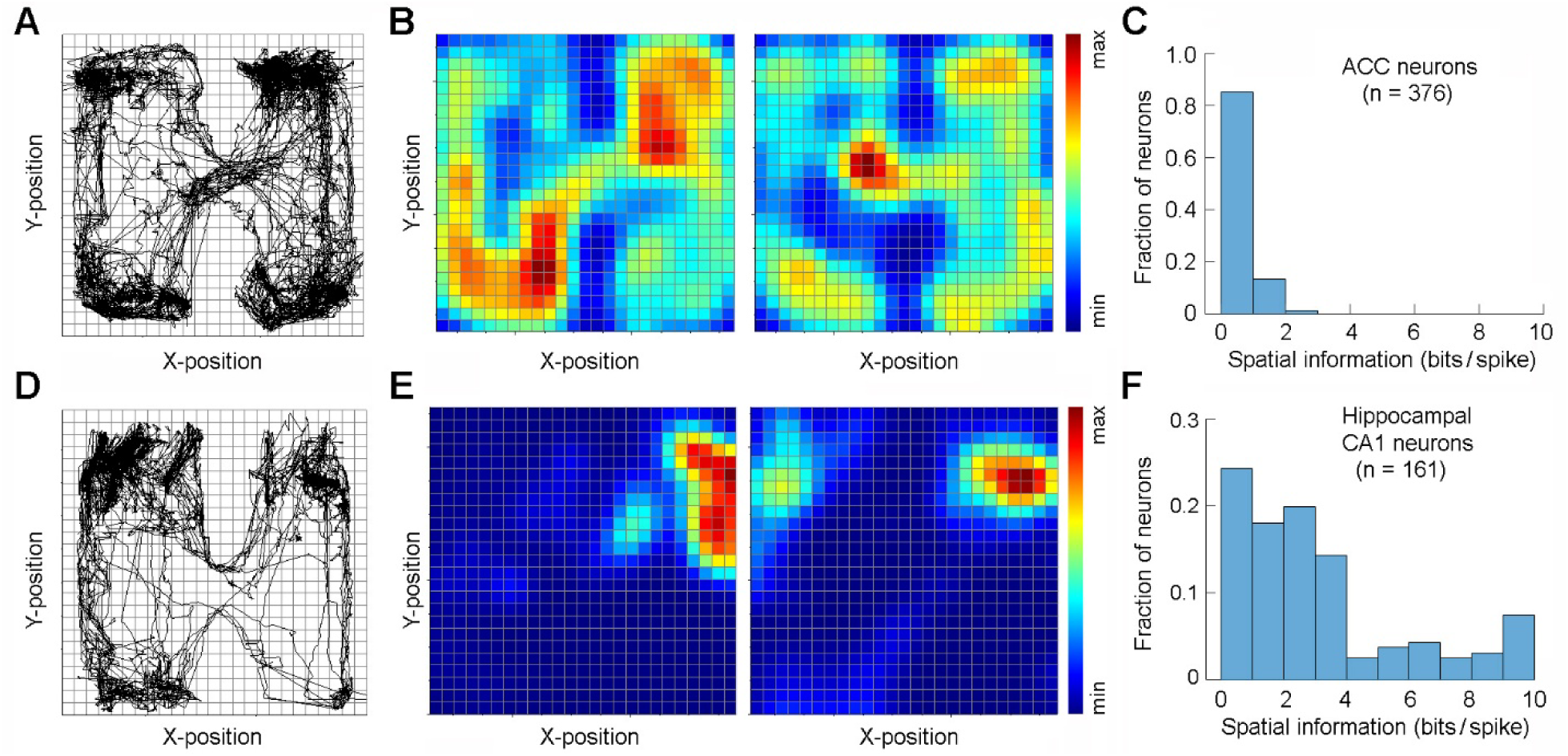
ACC neurons do not display place cell characteristics. **A,** A representative navigation path during inter-trial intervals of a discrimination–avoidance task session. **B,** Place field activity of two representative ACC neurons. Both neurons show spiking activity across the chamber without place preference. The color bar indicates normalized firing rate. **C,** Spatial information coding distribution of all recorded ACC neurons (n = 376). **D–F,** Same as in A–C, but for neurons recorded from hippocampal dorsal CA1 (n = 161). Notably, more than half of CA1 neurons exhibit high information coding (>2 bits/spike; F), whereas very few ACC neurons show comparable levels of information coding (C).

**Fig. S8.**
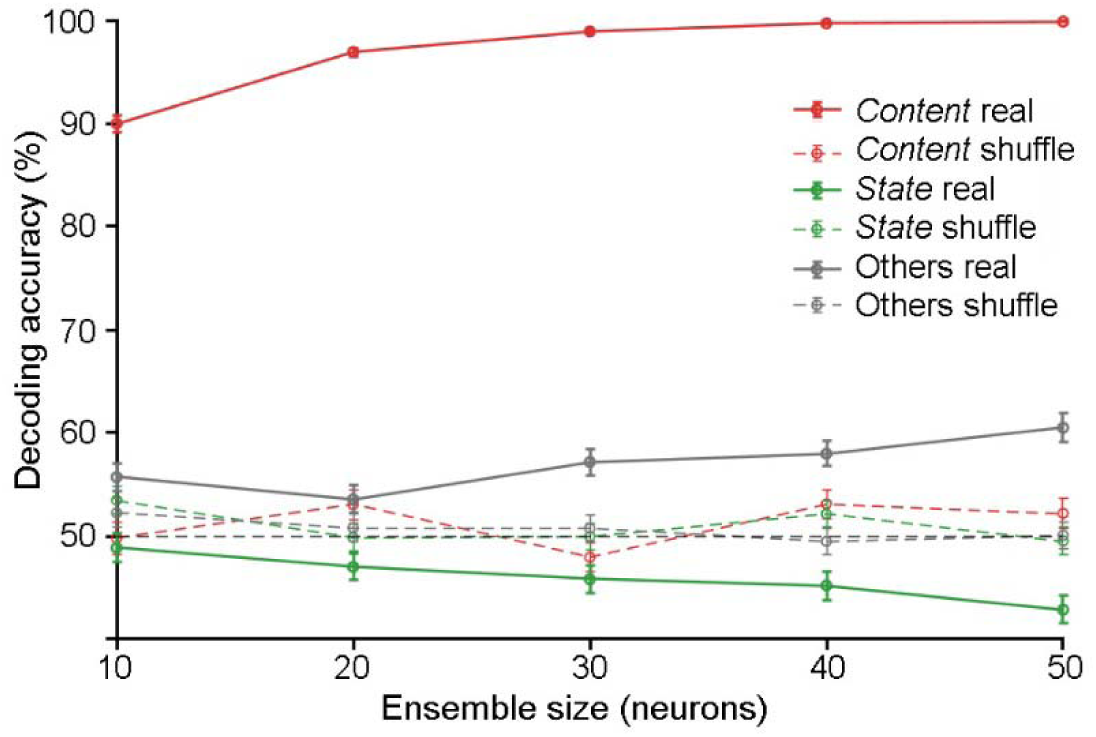
Pseudo-ensemble decoding dissociates content and state signals. Trial-level firing rates (0–5 s post-action window) were used to construct pseudo-population vectors from neurons classified as *action-content*, *action-state*, or other ACC neurons. For each ensemble size, neurons were randomly sampled and combined across sessions to generate pseudo-trials. The SVM was trained to classify action contents (A→B *vs.* B→A) and evaluated on held-out pseudo-trials. *Action-content* neurons supported robust decoding that scaled with ensemble size, whereas non-*action-content* neurons did not exceed shuffled controls, indicating that action content was selectively encoded in the content population. Shaded regions denote ±SEM across repetitions; dashed lines indicate label-shuffled controls.

**Fig. S9.**
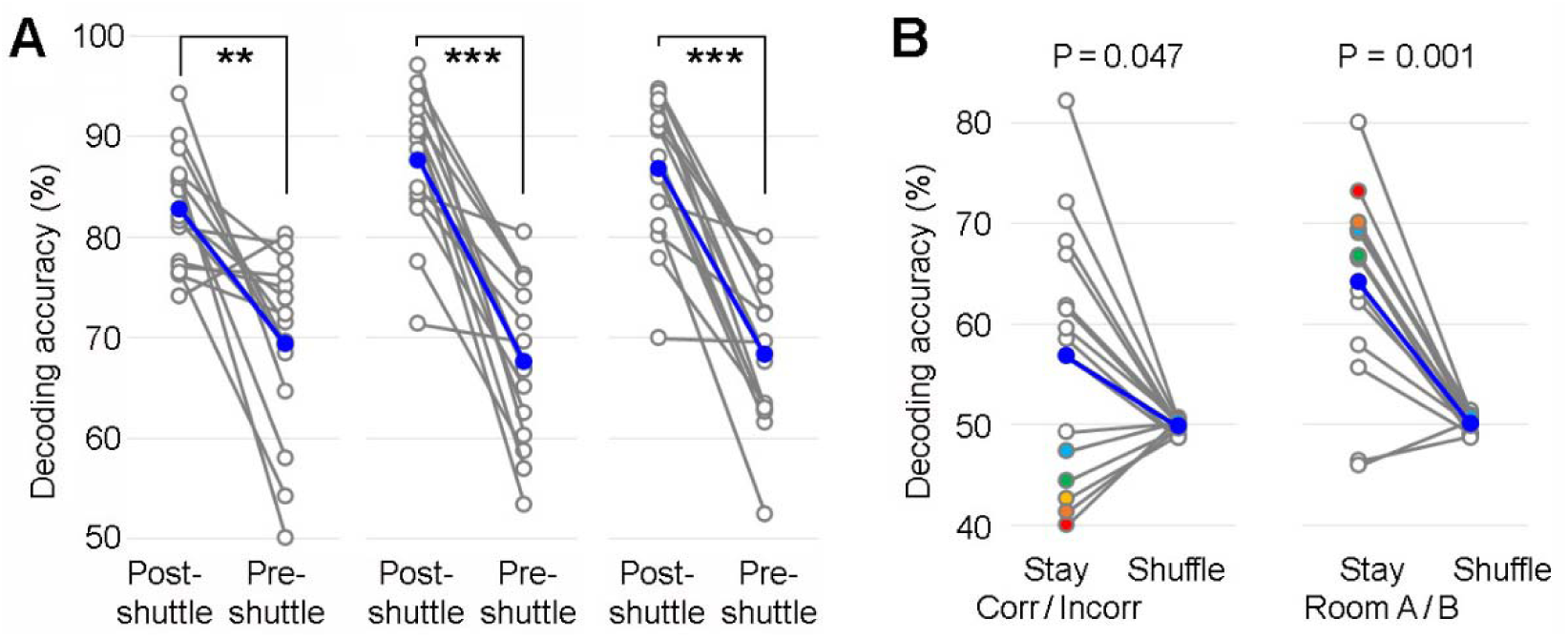
SVM decoding of multiple behaviors from ACC neuronal population activity. **A**, Mean (blue line) and individual session decoding accuracies (grey lines; 15 sessions) for decoding A→B *vs.* B→**A** shuttles using short (left, 2.5 s), medium (middle, 7.5 s), or long decoding windows (right, 10 s) surrounding shuttle crossings. **\*\***P < 0.01, **\*\*\***P < 0.001, *Wilcoxon signed-rank test*. **B**, Mean (blue line) and individual session decoding accuracies (grey lines) for decoding correct *vs.* incorrect stays (left) or Room-A *vs.* Room-B stays (right), using a 10-s window after sound onset. Notably, in several sessions, lower-than-chance decoding accuracy for Corr/Incorr stays (colored dots; left) corresponded to high decoding accuracy for Room A/B stays (shown in the same colors; right), suggesting a potential confound between these variables.

